# Explaining the unexplained admixture mapping signals via rare variants: the HCHS/SOL

**DOI:** 10.64898/2026.02.20.707089

**Authors:** Xueying Chen, Maria Argos, Bing Yu, Eric Boerwinkle, Qibin Qi, Robert Kaplan, Nora Franceschini, Tamar Sofer

**Affiliations:** Department of Biostatistics, Harvard T.H. Chan School of Public Health, Boston, MA, USA; CardioVascular Institute (CVI), Beth Israel Deaconess Medical Center, Boston, MA, USA; Department of Environmental Health, School of Public Health, Boston University, Boston, MA, USA; Division of Epidemiology and Biostatistics, School of Public Health, University of Illinois Chicago, Chicago, IL, USA; Department of Epidemiology, School of Public Health, The University of Texas Health Science Center at Houston, Houston, TX, USA; Department of Epidemiology & Population Health, Albert Einstein College of Medicine, Bronx, NY, USA; Division of Public Health Sciences, Fred Hutchinson Cancer Center, Seattle, WA, USA; Department of Genetics, University of North Carolina at Chapel Hill, Chapel Hill, NC, USA; Department of Epidemiology, University of North Carolina at Chapel Hill, Chapel Hill, NC, USA; Department of Medicine, Harvard Medical School, Boston, MA, USA

## Abstract

In admixed populations, formed by a mixing of two or more previously isolated populations, genomic segments can be traced to their ancestral populations (“ancestries”). Admixture mapping (AM) associates local ancestry with outcomes in admixed populations, detecting signals when causal variants differ in frequency or effect across ancestral populations. Prior work showed that adjusting for nearby GWAS-identified common variants does not fully explain some AM signals. Here, we assessed two approaches to explain the previously unexplained AM signal: (1) including sets of rare variants; (2) increasing the genomic region considered when searching for common variants. We studied these hypotheses comprehensively using a whole-genome sequencing dataset coupled with metabolomics from the Hispanic Community Health Study/Study of Latinos. We detected multiple sets of rare variants with replicated association with metabolite levels. Yet these rare variants appear to explain only a small fraction of the AM signal, while inclusion of common variants from a larger genomic region appears to explain the majority of the AM signals.

## Introduction

Admixture refers to the interbreeding of previously isolated ancestral populations, resulting in a new population in which genomic segments can be attributed to specific ancestral origins ^[1]^. Admixed individuals, for instance, those from Hispanic/Latino populations, possess genetic material from a combination of multiple ancestries (primarily European, African, and Amerindian ancestral populations). As a result, the presence of multiple ancestries contributes to unique linkage disequilibrium (LD) patterns both between and within Hispanic/Latino populations ^[1,2]^. Additionally, allele frequencies associated with disease risk or quantitative traits often differ across these ancestral groups. The underrepresentation of Hispanic/Latino individuals in genetic association studies ^[3,4]^, along with the complexities introduced by admixture, poses challenges for identifying causal variants and accurately estimating their effect sizes.

To address these challenges, some studies focusing on admixed populations leverage local ancestry information to identify potential variant associations with phenotype traits via admixture mapping (AM) ^[5–9]^. Local ancestry is defined at the variant or genomic segment level and refers to the reference ancestry from which the locus was inherited. At a given genomic position, the number of chromosomal copies with local ancestry from a given ancestry is coded (possibly 0, 1, or 2 copies) and referred as local ancestry counts, which is used in association analyses with the phenotype as the exposure measure ^[10]^. Local ancestry can be inferred through reference panels consisting of different populations ^[11,12]^. In particular, Fast Local Ancestry Inference Estimation (FLARE) is a recently developed method using an extended Li and Stephens model with high computational efficiency ^[13]^, and has been successfully applied to multiple admixed populations ^[14]^, including participants from the Hispanic Community Health Study/Study of Latinos (HCHS/SOL).

In AM, an association is observed when either the causal variant frequencies or effect sizes (or both) differ between the modeled ancestral populations ^[15,16]^. AM offers greater power than standard genome-wide association studies (GWAS) by potentially capturing more complex associations, such as rare variants and haplotypes, while also reducing the burden of multiple testing ^[16,17]^. However, the modeled local ancestry unit typically tags the causal variant through LD, which often extends over a broad genomic region ^[18]^. Due to the extensive length of genomic segments included in inferred local ancestries, a statistically significant signal from AM can cover many genetic variants, making pinpointing the divergence-driving variants challenging. A standard approach for identifying the causal variant is to align the AM results with the GWAS results ^[19]^. After finding the highly significant variants in GWAS, they can be adjusted in a regression and by checking the association with local ancestry counts, such that we can know which variants are driving the ancestral difference. However, this approach does not always successfully fully account for the observed association. A prior study using HCHS/SOL data conducted AM between local ancestry and metabolites, identifying genomic regions enriched for African and Amerindian ancestries that associated with metabolite levels. These regions were then mapped to relevant genes and common variants, the local ancestry-metabolite associations were adjusted to the suggested common variants, and sometimes the AM signal would become non-significant in this conditional analysis ^[19]^. The analysis was sometimes unable to identify specific genetic variants responsible for the observed associations—that is, common variants explaining the significant AM signals ^[19]^.

In GWAS, while common variants account for many phenotype associations ^[20]^, studying rare variants (RVs) remains essential. Recent advances in sequencing technology have made it possible to investigate rare variants with minor allele frequencies (MAF) < 0.01, which were previously difficult to detect using genotyping arrays and were imputed with lower accuracy than common variants ^[21,22]^. RVs have been reported to have important roles in complex traits and can exhibit large effect sizes ^[23–25]^, as some of the variants (e.g. putative loss-of-function [pLoF] and missense mutations) tend to be deleterious and occur at low allele frequencies ^[26,27]^.

However, the potential role of rare variants in relation to AM and local ancestries has not yet been thoroughly explored and understood. Therefore, in this study we investigated rare variants and their potential role in explaining the AM signal. We hypothesize that some of the unexplained AM-based associations with metabolites are due to rare variants that are potentially ancestry-enriched, i.e. are more likely to be on a genomic interval inherited from one of the parental ancestries in the admixed population. Identifying ancestry-enriched rare variants and their associations with metabolites can improve our understanding of the genetic basis of metabolic pathways and related diseases, especially in populations underrepresented in genetic studies. This approach may also aid in uncovering ancestry-specific risk factors and potential targets for drug development.

## Methods

### The HCHS/SOL

The Hispanic Community Health Study/Study of Latinos (HCHS/SOL) is a multi-center, population-based prospective cohort study established with participant enrollment between 2008 and 2011, with individuals of self-identified Hispanic/Latino background aged 18–74 years at baseline ^[28]^. The primary objective of the study is to investigate the prevalence and determinants of chronic conditions—including cardiovascular disease, asthma, and sleep disorders—within Hispanic/Latino populations ^[28]^. Participants were recruited from four U.S. field centers located in the Bronx (New York), Chicago (Illinois), San Diego (California), and Miami (Florida) ^[12,28,29]^, using a two-stage probability sampling design targeting households in census block groups enriched for Hispanic/Latino residents. Each study participant self-reported their Hispanic/Latino background, with major groups including Mexican, Puerto Rican, Cuban, Dominican, Central American, and South American ^[29,30]^. Baseline data collection included demographic information (sex and age at the time of participant’s clinic visit), clinical measurements such as body mass index (BMI), and estimated glomerular filtration rate (eGFR) ^[28]^. The eGFR was calculated from serum cystatin C concentrations using the Chronic Kidney Disease Epidemiology Collaboration equation, excluding demographic variables ^[31]^.

### Metabolomics data

Metabolomics analyses were conducted on fasting serum in two separate HCHS/SOL profiling efforts. In 2017, 4,002 HCHS/SOL participants who had also been genotyped were randomly selected for metabolomic profiling (“batch 1”). In 2021, an additional 2,368 serum samples from 2,330 participants (collected at baseline) were assayed (“batch 2”). Batch 2 contained repeated samples, including those from participants who had been analyzed in batch 1, for quality control. Batch 2 participants also included individuals who were sampled for various ancillary studies: it included (1) participants in the ECHO-SOL ancillary study of echocardiogram ^[32]^, (2) individuals whose eGFR was normal (i.e., > 60) at baseline but underwent a marked decline from the baseline to the second HCHS/SOL exam, and (3) individuals whose eGFR was measured at both the baseline and the second HCHS/SOL exams.

Serum samples were held at −70 °C in the HCHS/SOL Core Laboratory at the University of Minnesota until they were analyzed by Metabolon, Inc. (Durham, NC) in 2017 (batch 1) and 2021 (batch 2). Samples were then extracted and prepared using Metabolon’s standard solvent extraction protocol. The resulting extracts were divided into five fractions for use across four LC-MS–based metabolomic quantification platforms—two reverse-phase methods with positive-ion electrospray ionization (EI), one reverse-phase method with negative-ion EI, and one hydrophilic interaction liquid chromatography method with negative-ion EI—while the fifth fraction was reserved as a backup. Instrument variability was quantified by calculating the median relative standard deviation (SD) for internal standards added to each sample prior to mass spectrometry injection. Overall process variability was determined by computing the median relative SD for all endogenous metabolites (i.e., non-instrument standards) detected in 100% of the technical replicate samples.

Metabolites with proportion of missing values 0.25 or higher were excluded, and those with missing values with a proportion < 25% were imputed with the observed minimum value. All metabolite values were then inversely rank-normalized transformed to approximate a normal distribution, with rank randomly selected at ties, matching with the previous work ^[19]^. The same scaling and imputation procedure was performed separately across the two batches.

### Sequencing, quality control, and relatedness inference

The HCHS/SOL whole genome sequencing (WGS) data were generated at Baylor College of Medicine Human Genome Sequencing Center (HGSC) using the Trans-Omics for Precision Medicine (TOPMed) ^[33]^ libraries and protocols. Whole-genome sequencing (WGS) for TOPMed libraries was performed on NovaSeq 6000 instruments, generating 150 bp, dual-indexed, paired-end reads. Library pooling was conducted in two stages: an initial calibration pool to evaluate uniformity and perform quality control, followed by a re-pooling step to ensure a minimum sequencing depth of 30× per sample ^[34]^. Details about DNA sample handling, QC, as well as library construction, are publicly available. Then the DRAGEN pipeline was applied to the WGS data after QC ^[35]^.

We followed the approach used in previous studies ^[36]^ analyzing genetic associations in HCHS/SOL participants, incorporating genetic principal components and a kinship matrix. However, we generated these covariates using only the subset of individuals from the metabolite batches (n=5768), rather than the full WGS dataset. The detailed workflow is publicly available named TOPMed pipeline ^[37]^. We followed the procedure by using an iterative approach.

Variants from the WGS data were filtered first: SNPs that are bi-allelic, with MAF > 0.01, and missing rate < 0.01, and a pairwise r^2^ < 0.1 (r = 0.32) were selected. Furthermore, SNPs within regions such as lactase (LCT), human leukocyte antigen (HLA), or regions with polymorphic inversions on chromosomes 8 and 17 were excluded ^[36]^. We also excluded 19 individuals with outlying ancestry patterns, as identified by an initial PCA analysis. These individuals are those previously identified as having excess East Asian ancestry ^[36]^. We first computed the initial kinship matrix and population divergence estimates using KING robust ^[38]^ for individuals from both metabolomic batches combined, using the default minimum kinship of 2 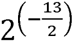 or 5th degree relatives as threshold. Then, PC-AiR ^[39]^ was used to compute genetic principal components (PCs), which selected an informative set of unrelated samples, computing PCA on them and project into relatives. We used 3^rd^ degree relatives as the kinship and ancestry divergence thresholds, with MAF and missingness thresholds of 0.01 and 0.05, respectively, to compute PCA correlations. After obtaining the initial PCs, PC-Relate was used to get the corrected kinship estimates ^[40]^. Then we followed another iteration of PC-AiR and PC-Relate to match the previous workflow ^[36]^, and eventually computed the corrected kinship matrix and genetic PCs.

### Local ancestry inference

FLARE v0.3.0 was used for local ancestry inference (LAI), as previously reported. HCHS/SOL individuals were genotyped using the Multi-Ethnic Genotyping Array (MEGA) ^[41]^, as part of their involvement in the Population Architecture Using Genomics and Epidemiology (PAGE) consortium ^[42]^. Genotypes were subsequently imputed to the TOPMed v1.0 reference panel ^[43]^.

Both the reference haplotypes and HCHS/SOL variant data were phased with Beagle v5.4 ^[44]^. SNPs were kept if they had a minor allele frequency ≥ 0.005 and, for imputed variants, an imputation quality score (R²) > 0.8, providing a stringent yet efficient quality threshold ^[13,45]^. All FLARE-derived coordinates are reported in GRCh38 (hg38). The reference haplotypes were generated from the Human Genome Diversity Project (HGDP) data ^[46,47]^, which were derived from whole-genome sequencing data of 220 participants across three geographic regions: Central-South America (61), Europe (155), and Sub-Saharan Africa (104). The FLARE-inferred local ancestry dataset comprises 11,928 HCHS/SOL individuals and 3,618,751 SNPs ^[41]^. Note that admixture mapping in this paper was performed on individuals with local ancestry information from the FLARE dataset who also had WGS data and were included in both metabolomic batches (n = 5307).

### Selected AM association regions

The previously published AM study identified statistically significant associations between metabolites and local Amerindian or African ancestry among participants in the HCHS/SOL cohort ^[19]^. To serve as negative controls, we selected four genomic regions where significant AM signals were found in association with metabolites (N2-acetyllysine, betaine, N-acetylglucosaminylasparagine, and cysteinylglycine) and could be attributed to common variants. In addition, we selected twelve test regions where the significant signals with the metabolites (N-acetylarginine, 3-aminoisobutyrate, 5-oxoproline, N-acetylputrescine, carnitine, 3beta-hydroxy- 5-cholestenoate, arachidonoyl choline, 1-methylimidazoleacetate, ethylmalonate, N-acetylcarnosine, propyl 4-hydroxybenzoate sulfate, and cys-gly, oxidized) were not explained by common variants.

### The modified STAAR pipeline

For detecting potential rare variant associations, we adapted the STAAR pipeline ^[48,49]^ to identify rare variant sets that might explain these unexplained AM signals. The modified STAAR pipeline is illustrated in Figure 1, showing how associated common and rare variant sets were identified and incorporated into downstream analysis, using specific genomic intervals rather than genome-wide testing. The metabolite-associated AM regions were identified in the previous work on the level of genomic intervals ^[19]^. We defined genomic association regions with AM signal as all consecutive local ancestry intervals with genome-wide significant AM associations with the metabolite of interest (potentially including multiple adjacent intervals in addition to the most significant interval) (Table 1). For AM signals that only span one local ancestry interval, we expanded the interval along with a ±1 megabase window for analysis, or until reaching the start or end of the chromosome, whichever occurred first. Sets of rare variants (MAF < 0.01) were then constructed through the adapted pipeline ^[4]^. We performed gene-centric rare variant set associations for coding and non-coding functional categories (Figure 1). STAAR employs burden tests, variance component tests, with a multi-dimensional omnibus weighting framework to incorporate various variant-annotation scores and qualitative functional categories for genetic variants, where the result of different tests were reported but we only focused on the STAAR burden (STAAR-B) and the STAAR omnibus (STAAR-O) test result to evaluate whether or not a rare variant set has significant association with the metabolite. We modified the STAAR pipeline to extract burden scores, which could be combined with the local ancestry association in a conditional model. Under the framework of using conditional analysis with both local ancestries and rare variants in the same model to establish that the rare variants explain the AM signal, we need to extract an effect size of the local ancestry dosage while adjusting for the rare variant set. This is impossible when using a variance component test such as SKAT, as it does not allow for estimation of the impact of the variant set on the AM-metabolite association. Therefore, we used the weights from the burden tests in STAAR to generate Burden scores, which were then used as covariates in the conditional admixture mapping analysis.

**Figure 1.**
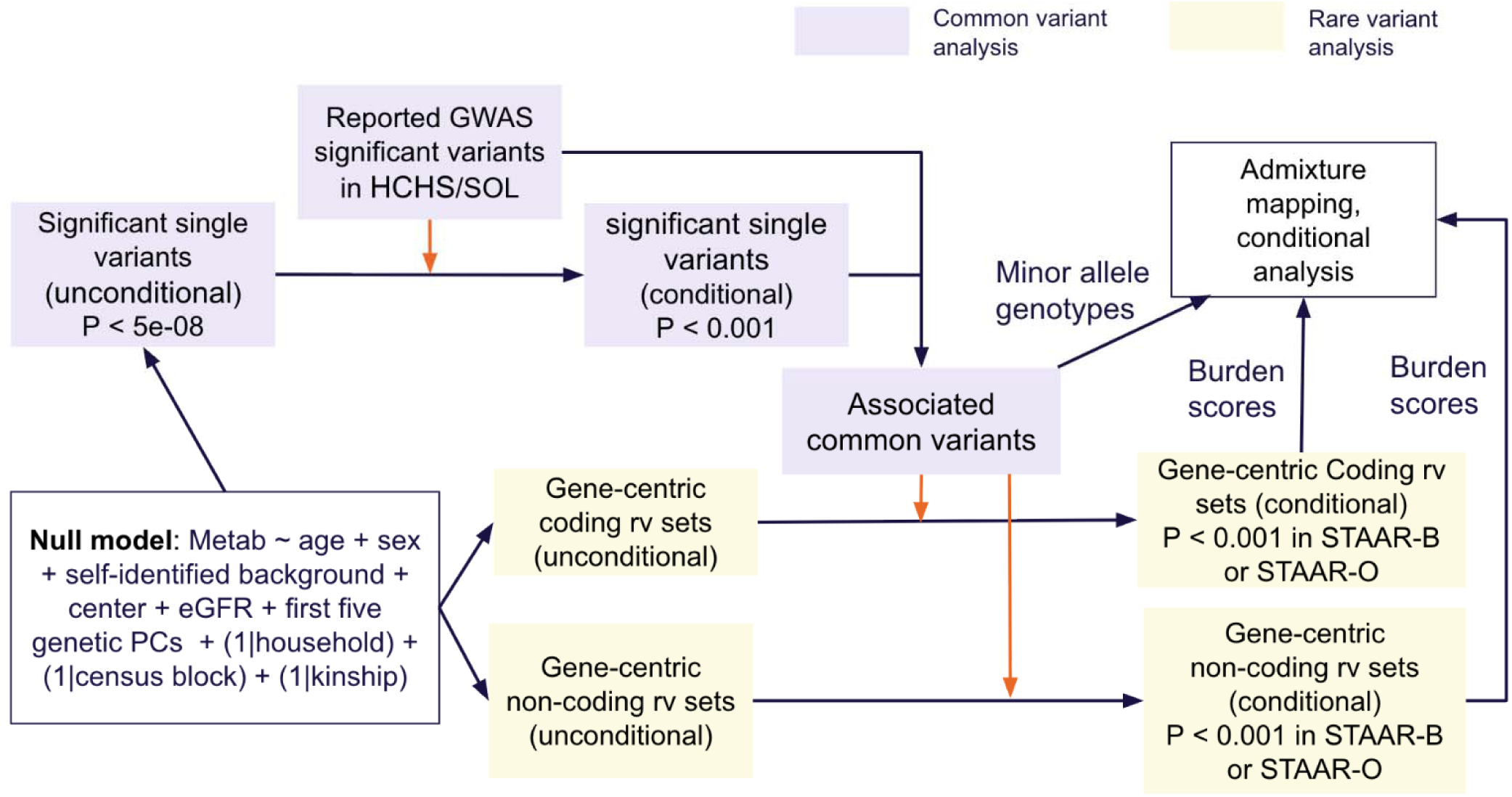
Workflow for identifying common and/or rare variant sets explaining an AM signal using the adapted STAAR pipeline. First starting with fitting the null model: metabolite value as outcome, adjusting covariates shown above and random effects of household, census block, and kinship. Next steps are color-coded by variant type: lavender for common variant analysis and yellow for rare variant analysis. Orange arrows indicate conditional analyses that adjust for the variants or sets in the target box. The significance thresholds for gene-centric unconditional analyses (both coding and non-coding) were computed by P < 0.05/(the number of genes x the number of function categories). For all conditional variant associations, a nominal p-value (0.001) was used. Conditional admixture mapping was performed using four models per metabolite–region pair: null, common variants only, rare variant burden only, and a combined model including both. PC: principal component; eGFR: estimated glomerular filtration rate; AM: admixture mapping; RV: rare variants; STAAR-B: STAAR Burden p-value; STAAR-O: STAAR omnibus p-value.

**Table 1.**
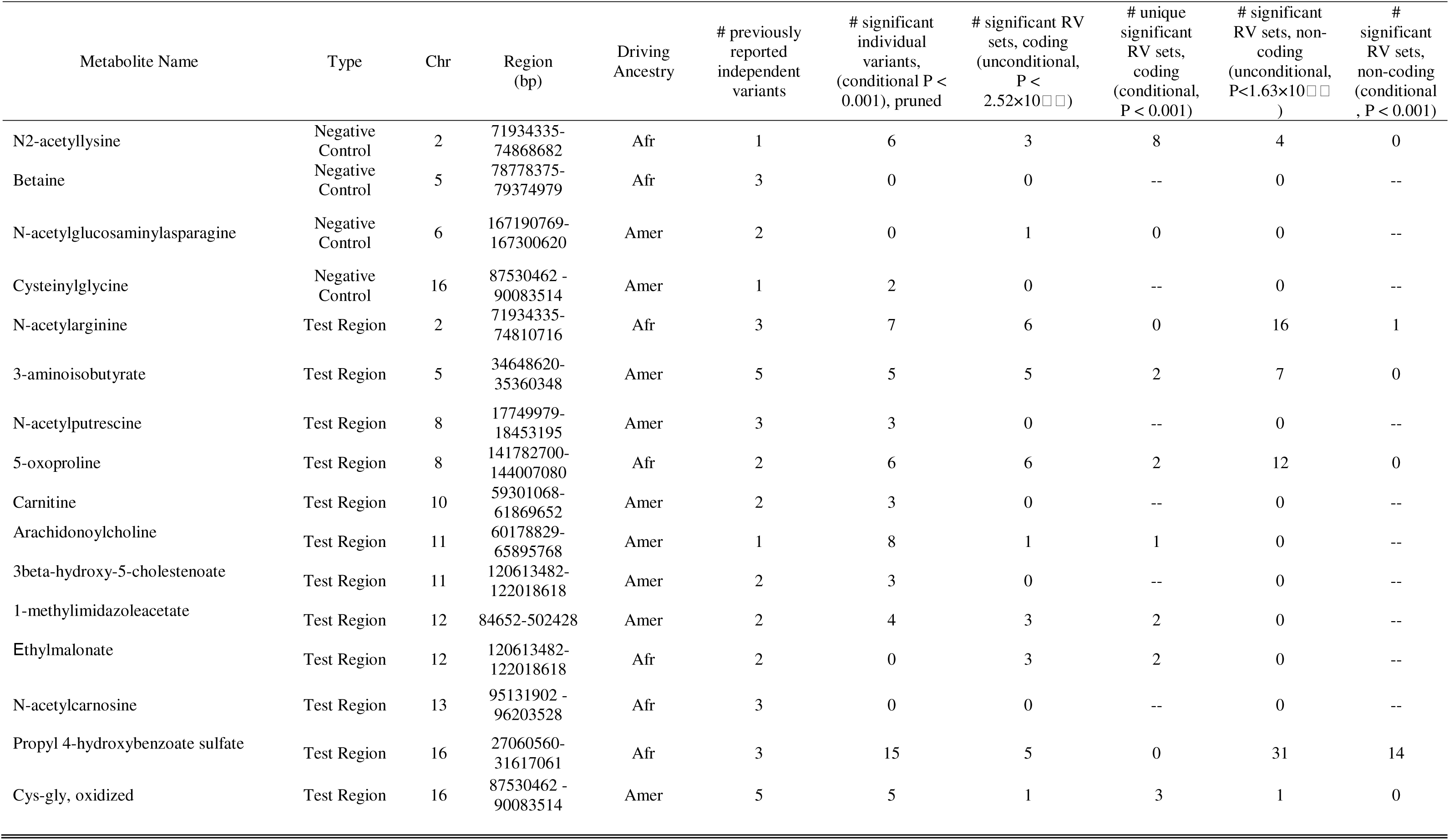
Number of significant single common variants and rare variant sets identified using the modified STAAR pipeline, both conditional and unconditional on the “associated common variants” for each trait and genomic region, which were generated by combining the variants from the 6^th^ and 7^th^ column in the table together. Afr: African ancestry, Amer: Amerindian ancestry, RV: rare variants.

| Metabolite Name | Type | Chr | Region (bp) | Driving Ancestry | # previously reported independent variants | # significant individual variants, (conditional P < 0.001), pruned | # significant RV sets, coding (unconditional, P < 2.52×10<sup>-6</sup>) | # unique significant RV sets, coding (conditional, P < 0.001) | # significant RV sets, non-coding (unconditional, P < 1.63×10<sup>-6</sup>) | # significant RV sets, non-coding (conditional, P < 0.001) |
| --- | --- | --- | --- | --- | --- | --- | --- | --- | --- | --- |
| N2-acetyllysine | Negative Control | 2 | 71934335-74868682 | Afr | 1 | 6 | 3 | 8 | 4 | 0 |
| Betaine | Negative Control | 5 | 78778375-79374979 | Afr | 3 | 0 | 0 | -- | 0 | -- |
| N-acetylglucosaminylasparagine | Negative Control | 6 | 167190769-167300620 | Amer | 2 | 0 | 1 | 0 | 0 | -- |
| Cysteinylglycine | Negative Control | 16 | 87530462 - 90083514 | Amer | 1 | 2 | 0 | -- | 0 | -- |
| N-acetylarginine | Test Region | 2 | 71934335-74810716 | Afr | 3 | 7 | 6 | 0 | 16 | 1 |
| 3-aminoisobutyrate | Test Region | 5 | 34648620-35360348 | Amer | 5 | 5 | 5 | 2 | 7 | 0 |
| N-acetylputrescine | Test Region | 8 | 17749979-18453195 | Amer | 3 | 3 | 0 | -- | 0 | -- |
| 5-oxoproline | Test Region | 8 | 141782700-144007080 | Afr | 2 | 6 | 6 | 2 | 12 | 0 |
| Carnitine | Test Region | 10 | 59301068-61869652 | Amer | 2 | 3 | 0 | -- | 0 | -- |
| Arachidonoylcholine | Test Region | 11 | 60178829-65895768 | Amer | 1 | 8 | 1 | 1 | 0 | -- |
| 3beta-hydroxy-5-cholestenoate | Test Region | 11 | 120613482-122018618 | Amer | 2 | 3 | 0 | -- | 0 | -- |
| 1-methylimidazoleacetate | Test Region | 12 | 84652-502428 | Amer | 2 | 4 | 3 | 2 | 0 | -- |
| Ethylmalonate | Test Region | 12 | 120613482-122018618 | Afr | 2 | 0 | 3 | 2 | 0 | -- |
| N-acetylcarnosine | Test Region | 13 | 95131902 - 96203528 | Afr | 3 | 0 | 0 | -- | 0 | -- |
| Propyl 4-hydroxybenzoate sulfate | Test Region | 16 | 27060560-31617061 | Afr | 3 | 15 | 5 | 0 | 31 | 14 |
| Cys-gly, oxidized | Test Region | 16 | 87530462 - 90083514 | Amer | 5 | 5 | 1 | 3 | 1 | 0 |

### Common variant associations

For each metabolite, we first fit a linear mixed “null” model, i.e. without including any of the variants of interest, adjusting for age, sex, the first 5 genetic principal components, self-reported backgrounds, eGFR, and incorporating random effects for household, census block, and genetic relatedness (kinship matrix).

In an attempt to expand the common variants selected to explain the AM signal in the defined genomic regions, we first performed single-variant association tests in batch 1 to identify significantly associated common variants (P < 5×10□□). We then performed conditional analyses by adjusting for common variants previously associated with each metabolite in HCHS/SOL, as reported in the GWAS Catalog ^[20]^, to identify additional associated common variants (P < 0.001) that are independent of the reported GWAS hits. LD-pruning was performed for the additional set of common variants using a sequential conditional analysis based on the null model via score tests ^[48]^. Following pruning, the additional associated variants were combined with the catalog-reported common variants to generate the final set of “associated common variants” for each metabolite. This set was then used in subsequent conditional analyses of rare variants and in conditional AM (Figure 1). We also calculated the span (maximum distance in base pairs) of the “associated common variants” within each genomic region.

### Rare variants testing in the modified STAAR pipeline

Rare variant association analysis was conducted separately for the two batches, using all individuals with available WGS and local ancestry data—3842 in batch 1. STAAR pipeline uses multiple burden tests for each set of rare variants, with weights based on ranks from functional annotations, as well as on variant frequencies via the beta distributions with different parameters, to upweight rare variants ^[49]^. Protein-coding genes were selected by Ensembl and non-coding RNA (ncRNA) genes selected by GENCODE ^[48]^. The coding functional categories include plof (putative loss of function), plof_ds (putative loss of function or disruptive missense), missense, disruptive missense, synonymous, ptv (protein-truncating variant), and ptv_ds (protein-truncating variants or disruptive missense). The non-coding categories include downstream, upstream, UTR (untranslated region), CAGE (cap analysis of gene expression)-defined promoters and enhancers, DHS (DNase I hypersensitive site) defined promoters and enhancers, and non-coding RNAs (ncRNA).

Using the modified pipeline, we extracted all burden scores from STAAR default burden tests, and evaluated their correlations with each other, then selected a smallest set of burden score that is not highly correlated (r < 0.85), because the burden scores generated by two Beta distributions were highly correlated, one of burden scores under either the Beta (1,1) or Beta (1, 25) weighting could be served as the final covariate fitting into the conditional AM model (Figure 1).

We used Bonferroni correction to defined statistically significant burden scores in unconditional rare variant association analysis calculated separately for coding (P < 2.52×10 for batch 1 and P < 2.71×10□□ for batch 2) and non-coding (P < 1.63×10□□ for batch 1 and P < 1.47×10□□ for batch 2) analyses based on the number of tests performed in each gene and each functional category. For discovering rare variant sets, we also reported SKAT and the STAAR-O omnibus p-values. However, a variant set is considered statistically significant in association with the metabolite only if its p-value is below the computed thresholds for any of the STAAR-O or STAAR-B (1, 25) omnibus p-values.

### Conditional admixture mapping with identified variants

After identifying known common variants and obtaining the burden scores from the rare variant sets, we performed conditional analyses by re-fitting the AM model in four ways: first without any genetic variants, only with the “associated common variants”, only with the rare variant sets, and finally with both the known common variants and the rare variant sets. This approach allowed us to see whether the local ancestry associations became statistically insignificant, thus revealing if these variants accounted for the observed associations. The AM conditional analyses are then conducted in the replication (batch 2, N = 1465) dataset for validation. The significance threshold in AM using FLARE local ancestries was previously estimated using STEAM ^[41]^, a method that quantifies the hypothesis testing burden in AM based on ancestry correlation structure and the number of generations since admixture ^[50]^.

## Results

### Participant characteristics

Participant characteristics for those included in the metabolomics analysis are summarized in Supplementary Table S1, stratified by metabolomics batch. The average age was approximately 46 years in the discovery batch (batch 1) and 53 years in the replication batch (batch 2).

Compared to batch 1, batch 2 included a higher proportion of female participants (∼63% vs. ∼57%), a greater prevalence of hypertension and/or diabetes, and lower average eGFR levels. The distribution of participants across recruitment centers was similar between the two batches (Supplementary Table S1).

Principal components (PCs) were computed using combined data from both metabolomics batches, and the first five PCs were found to capture most of the genetic variation and effectively account for population substructure (Supplementary Figure S1). Notably, starting from PC5 and extending through PC32, a clustering pattern emerged among self-identified Mexican individuals (Supplementary Figure S1), which may reflect the influence of small sample sizes and the use of whole genome sequencing (WGS) data, capturing fine-scale population structure. Overall, the separation across different self-identified backgrounds was minimal between PC5 and PC6, and using the first five PCs is consistent with previous findings and established methodologies.

### Single variant and rare variant sets identified by the modified STAAR pipeline

Following the workflow (Figure 1), for each metabolite and association region of interest we defined a set of “associated common variants” by identifying GWAS-significant variants, using previously reported GWAS-significant variants, and next by performing conditional analysis with these two sets of variants adjusted for (Figure 1).

In the gene-centric coding and non-coding analysis, we identified significant rare variant sets, for any of the STAAR-O, STAAR-B (1,1), or STAAR-B (1, 25) p-value, in two negative control regions and two test regions in the unconditional analysis (Table 1, Supplementary Table S2). Table 1 summarized the number of significant rare variant sets detected using the modified STAAR pipeline, both before and after conditioning on the associated common variants in batch 1. After adjusting the associated common variants, rare variant sets in 9 out of 16 regions persisted (Table 1). Among rare variant sets sharing the same functional category within the same genomic region, 4 of the 9 regions showed replication of one or more rare variant sets (Supplementary Table S2).

Rare variant sets identified in coding regions were observed in 7 of the 9 regions (Tables 1 and 2). Notably, *AGXT2* on chromosome 5 associated with 3-aminoisobutyrate, *OPLAH* on chromosome 8 associated with 5-oxoproline, and *ACADS* on chromosome 12 associated with ethylmalonate exhibited concordant rare variant sets with identical functional categories, predominantly disruptive missense variants and predicted loss-of-function variants, optionally weighted by a deleteriousness score (plof_ds) (Table 2; Supplementary Table S3).

**Table 2.** Significant rare variant sets and their associated functional categories identified using the modified STAAR pipeline, conditioned on previously associated common variants, for each metabolite and genomic region. The reported p-values are omnibus p-values derived from multiple rare variant tests (SKAT, Burden, and ACAT), with the final integrated p-value summarized as STAAR-O. STAAR-S: the omnibus p-value for all SKAT tests under the specified Beta distribution. STAAR-B: the omnibus p-value for all Burden tests under the specified Beta distribution. STAAR-A: the omnibus p-value for all ACAT-V tests under the specified Beta distribution. ACAT-O: the omnibus p-value for all aggregated tests including the SKAT, Burden, and ACAT-V tests across two Beta distributions using Cauchy method. STAAR-O: the omnibus p-value for all aggregated tests including the SKAT, Burden, and ACAT-V tests weighted by annotation using Cauchy method.

| Gene Name | Chr | Metabolite | Group | Category | #SNV | cMAC | STAAR-S<br>(1,25) | STAAR-S<br>(1,1) | STAAR-B<br>(1,25) | STAAR-B<br>(1,1) | STAAR-A<br>(1,25) | STAAR-A<br>(1,1) | ACAT-O | STAAR-O |
| --- | --- | --- | --- | --- | --- | --- | --- | --- | --- | --- | --- | --- | --- | --- |
| AUP1 | 2 | N2-acetyllysine | Negative Control | plof_ds | 10 | 63 | 1.48E-03 | 1.46E-03 | 7.84E-04 | 6.74E-04 | 1.38E-03 | 1.37E-03 | 1.02E-03 | 1.08E-03 |
| CYP26B1 | 2 | N2-acetyllysine | Negative Control | missense | 31 | 135 | 2.06E-02 | 3.61E-02 | 1.41E-02 | 1.59E-02 | 1.80E-04 | 2.03E-04 | 5.33E-04 | 5.62E-04 |
| ALMS1 | 2 | N2-acetyllysine | Negative Control | missense | 238 | 974 | 1.38E-08 | 1.17E-07 | 2.35E-01 | 1.38E-01 | 1.47E-04 | 1.51E-04 | 2.17E-07 | 7.41E-08 |
| NAT8 | 2 | N2-acetyllysine | Negative Control | missense_cond | 22 | 56.06 | 3.90E-05 | 3.83E-05 | 1.69E-03 | 1.27E-03 | 3.51E-05 | 3.48E-05 | 5.43E-05 | 5.43E-05 |
| AUP1 | 2 | N2-acetyllysine | Negative Control | disruptive_missense_cond | 9 | 62 | 1.47E-03 | 1.45E-03 | 9.12E-04 | 7.81E-04 | 1.39E-03 | 1.37E-03 | 1.10E-03 | 1.15E-03 |
| EGR4 | 2 | N2-acetyllysine | Negative Control | synonymous_cond | 18 | 81 | 4.39E-06 | 4.30E-06 | 1.10E-03 | 5.76E-04 | 3.97E-06 | 3.92E-06 | 6.21E-06 | 6.19E-06 |
| ALMS1 | 2 | N2-acetyllysine | Negative Control | synonymous_cond | 81 | 603 | 1.36E-06 | 2.45E-06 | 9.00E-01 | 8.10E-01 | 7.94E-04 | 6.82E-04 | 6.81E-06 | 5.24E-06 |
| STAMBP | 2 | N2-acetyllysine | Negative Control | synonymous | 7 | 82 | 1.13E-03 | 1.28E-03 | 1.69E-04 | 1.60E-04 | 1.86E-03 | 1.82E-03 | 3.80E-04 | 4.03E-04 |
| DUSP11 | 2 | N-acetylarginine | Test Regions | enhancer_CAGE | 10 | 16 | 4.42E-05 | 4.31E-05 | 3.00E-01 | 2.90E-01 | 3.00E-01 | 2.90E-01 | 1.33E-04 | 1.31E-04 |
| AGXT2 | 5 | 3-aminoisobutyrate | Test Regions | plof_ds | 15 | 58 | 5.30E-06 | 6.51E-06 | 8.40E-11 | 1.10E-10 | 8.45E-06 | 8.98E-06 | 1.57E-10 | 2.86E-10 |
| AGXT2 | 5 | 3-aminoisobutyrate | Test Regions | disruptive_missense | 11 | 52 | 4.54E-06 | 5.74E-06 | 7.37E-11 | 9.82E-11 | 5.14E-06 | 5.73E-06 | 1.85E-10 | 2.53E-10 |
| OPLAH | 8 | 5-oxoproline | Test Regions | plof_ds | 14 | 23 | 1.22E-02 | 1.25E-02 | 3.83E-05 | 3.91E-05 | 3.83E-05 | 3.91E-05 | 5.09E-05 | 5.79E-05 |
| OPLAH | 8 | 5-oxoproline | Test Regions | disruptive_missense | 11 | 19 | 9.19E-03 | 9.43E-03 | 1.12E-04 | 1.14E-04 | 1.12E-04 | 1.14E-04 | 1.64E-04 | 1.69E-04 |
| MS4A14 | 11 | Arachidonoylcholine | Test Regions | synonymous | 25 | 175 | 4.96E-03 | 6.28E-03 | 1.24E-04 | 1.42E-04 | 1.51E-02 | 1.43E-02 | 3.57E-04 | 3.85E-04 |
| SLC6A13 | 12 | 1-methylimidazoleacetate | Test Regions | plof_ds | 20 | 71 | 1.12E-04 | 1.28E-04 | 1.07E-05 | 8.90E-06 | 7.01E-04 | 6.77E-04 | 2.59E-05 | 2.66E-05 |
| SLC6A13 | 12 | 1-methylimidazoleacetate | Test Regions | disruptive_missense | 18 | 69 | 1.13E-04 | 1.31E-04 | 5.17E-06 | 4.41E-06 | 7.00E-04 | 6.77E-04 | 1.28E-05 | 1.37E-05 |
| ACADS | 12 | Ethylmalonate | Test Regions | plof_ds | 26 | 34 | 7.56E-08 | 7.75E-08 | 6.26E-10 | 6.09E-10 | 6.26E-10 | 6.09E-10 | 3.37E-09 | 9.23E-10 |
| ACADS | 12 | Ethylmalonate | Test Regions | disruptive_missense | 25 | 33 | 6.26E-08 | 6.42E-08 | 3.02E-10 | 2.95E-10 | 3.02E-10 | 2.95E-10 | 9.09E-10 | 4.47E-10 |
| FAM57B | 16 | Propyl 4-hydroxybenzoate sulfate | Test Regions | enhancer_DHS | 66 | 288 | 2.60E-03 | 3.12E-03 | 5.96E-05 | 4.45E-05 | 3.33E-02 | 3.28E-02 | 1.17E-04 | 1.50E-04 |
| FUS | 16 | Propyl 4-hydroxybenzoate sulfate | Test Regions | enhancer_DHS | 123 | 778 | 3.51E-02 | 3.77E-02 | 2.79E-04 | 1.78E-04 | 1.22E-01 | 1.19E-01 | 6.31E-04 | 6.48E-04 |
| IL4R | 16 | Propyl 4-hydroxybenzoate sulfate | Test Regions | UTR | 55 | 425 | 4.24E-05 | 2.20E-05 | 2.50E-03 | 1.12E-03 | 6.00E-05 | 4.98E-05 | 5.70E-05 | 5.61E-05 |
| IL4R | 16 | Propyl 4-hydroxybenzoate sulfate | Test Regions | promoter_CAGE | 33 | 162 | 3.01E-06 | 2.94E-06 | 7.74E-04 | 3.04E-04 | 5.25E-06 | 4.77E-06 | 5.26E-06 | 5.57E-06 |
| IL4R | 16 | Propyl 4-hydroxybenzoate sulfate | Test Regions | promoter_DHS | 84 | 663 | 3.81E-07 | 4.30E-07 | 2.64E-04 | 1.42E-04 | 8.84E-05 | 7.68E-05 | 7.35E-07 | 1.20E-06 |
| IL4R | 16 | Propyl 4-hydroxybenzoate sulfate | Test Regions | enhancer_CAGE | 92 | 479 | 4.53E-05 | 3.53E-05 | 1.05E-02 | 8.30E-03 | 5.85E-05 | 5.01E-05 | 6.05E-05 | 6.84E-05 |
| IL4R | 16 | Propyl 4-hydroxybenzoate sulfate | Test Regions | enhancer_DHS | 507 | 3001 | 1.73E-03 | 6.48E-04 | 6.17E-04 | 4.83E-04 | 3.01E-04 | 2.52E-04 | 5.12E-04 | 4.58E-04 |
| ITGAL | 16 | Propyl 4-hydroxybenzoate sulfate | Test Regions | promoter_DHS | 54 | 145 | 2.78E-03 | 2.95E-03 | 6.65E-04 | 5.82E-04 | 2.39E-02 | 2.49E-02 | 1.63E-03 | 1.50E-03 |
| ITGAL | 16 | Propyl 4-hydroxybenzoate sulfate | Test Regions | enhancer_DHS | 63 | 261 | 5.72E-02 | 6.59E-02 | 1.27E-04 | 1.34E-04 | 7.60E-02 | 8.20E-02 | 4.01E-04 | 3.90E-04 |
| KDM8 | 16 | Propyl 4-hydroxybenzoate sulfate | Test Regions | enhancer_DHS | 194 | 1078 | 1.76E-04 | 2.48E-04 | 4.52E-02 | 2.77E-02 | 2.14E-02 | 2.03E-02 | 9.76E-04 | 6.09E-04 |
| MVP | 16 | Propyl 4-hydroxybenzoate sulfate | Test Regions | UTR | 26 | 115 | 4.11E-04 | 4.42E-04 | 8.42E-02 | 4.35E-02 | 5.21E-04 | 5.15E-04 | 7.03E-04 | 6.99E-04 |
| NUPR1 | 16 | Propyl 4-hydroxybenzoate sulfate | Test Regions | promoter_DHS | 6 | 43 | 5.41E-04 | 5.48E-04 | 9.14E-03 | 7.47E-03 | 6.88E-04 | 6.77E-04 | 8.71E-04 | 8.76E-04 |
| PRRT2 | 16 | Propyl 4-hydroxybenzoate sulfate | Test Regions | enhancer_DHS | 294 | 1078 | 1.46E-05 | 3.88E-05 | 9.36E-03 | 6.37E-03 | 4.30E-03 | 4.32E-03 | 6.13E-05 | 6.33E-05 |
| RNF40 | 16 | Propyl 4-hydroxybenzoate sulfate | Test Regions | UTR | 31 | 119 | 7.45E-05 | 9.75E-05 | 3.56E-03 | 3.23E-03 | 2.15E-03 | 2.15E-03 | 1.63E-04 | 2.38E-04 |
| DPEP1 | 16 | Cys-gly, oxidized | Test Regions | plof | 3 | 4 | 5.76E-03 | 5.80E-03 | 3.73E-04 | 3.74E-04 | 3.73E-04 | 3.74E-04 | 4.28E-04 | 5.43E-04 |
| DPEP1 | 16 | Cys-gly, oxidized | Test Regions | plof_ds | 6 | 10 | 6.90E-04 | 7.19E-04 | 5.38E-07 | 5.47E-07 | 5.38E-07 | 5.47E-07 | 5.88E-07 | 8.13E-07 |
| DPEP1 | 16 | Cys-gly, oxidized | Test Regions | disruptive_missense | 3 | 6 | 5.24E-03 | 5.33E-03 | 3.77E-04 | 3.80E-04 | 3.77E-04 | 3.80E-04 | 5.15E-04 | 5.48E-04 |

From the non-coding analysis, one genomic region contained significant rare variant sets that also have replicated results in batch 2 after conditioning on the associated common variants (Supplementary Table S3). In particular, the region at chromosome 16 associated with propyl 4-hydroxybenzoate sulfate included three replicated rare variant sets, including *FAM57B* and *PRRT2* (enhancer region that overlaps DHS site), and *RNF40* (UTR region). The limited number of rare variant sets identified after conditioning on common variants suggests that, for all 16 selected genomic regions, most of the genetic associations were largely accounted for by the common variants.

### Admixture mapping with rare variants – negative controls

After adjusted for the “associated common variants”, the AM signals were substantially reduced across the four negative control regions, and became statistically insignificant (Figure 2, Table 3). In batch 2, three out of four AM signals displayed patterns consistent with those observed in batch 1; however, the African local ancestry association with betaine did not replicate in batch 2, even without conditioning on any common variant genotypes (Figure 2). As expected, the AM signals were similarly explained across both batch 1 and batch 2, except for the local African ancestry association with betaine on chromosome 5 (Supplementary Figure S3, Table S2). We identified eight significant rare variant sets in one of the negative controls on chromosome 2 in batch 1, spanning multiple genes and functional categories: missense or disruptive missense variants (*AUP1, ALMS1, NAT8, CYP26B1*), plof_ds (*AUP1*), and synonymous variants (*EGR4, ALMS1, STAMBP*) (Table 2). However, conditional admixture mapping incorporating the corresponding burden scores resulted in only a partial reduction of the admixture signal, which remained above the significance threshold (Figure 2; Table 3). None of the rare variant sets mentioned above were replicated in batch 2 (Supplementary Table S2, S3).

**Figure 2.**
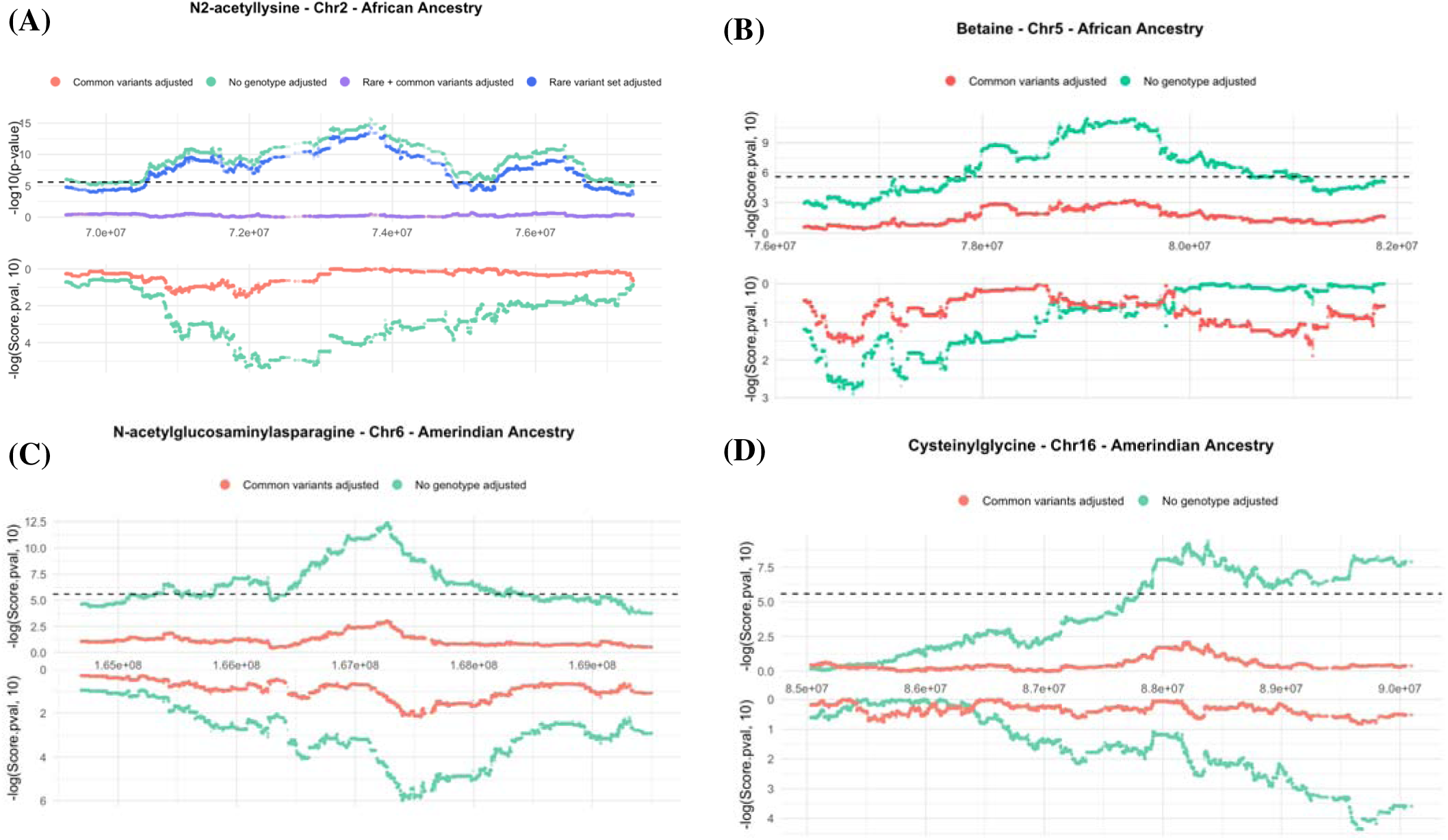
AM results in terms of -log10(P) using the driving local ancestries for four “negative control” associations, defined as those with associations explained by common variants or rare variant sets (if identified): (A) N2-acetyllisine on chromosome 2, (B) betaine on chromosome 5 (both driven by African ancestry), (C) N-acetylglucosaminylasparagine on chromosome 6, and (D) cysteinylglycine on chromosome 16 (both driven by Amerindian ancestry). For each trait, mirrored plots display results from the discovery dataset (batch 1, upper panel) and the replication dataset (batch 2, lower panel). The horizontal dashed black line in the upper panel denotes the AM significance threshold (P = 2.58 × 10^-6^).

**Table 3.** AM results for all 16 metabolite–genomic region pairs are shown, including effect size estimates and p-values of the lead AM variant under four models: the null model (no genotype adjustment), adjustment for associated common variants only, adjustment for rare variant sets using burden scores, and adjustment for both common and rare variants (if a rare variant set was identified).

| Metabolite Name | Group | Chr | Region (bp) | Driving Ancestry | AM lead variant Est. null model | AM lead variant p-value null model | AM lead variant Est. adjusting associated common variants | AM lead variant p-value adjusting associated common variants | AM lead variant Est. adjusting rv set only | AM lead variant p-value adjusting rv set only | AM lead variant Est. adjusting rare + associated common variants | AM lead variant p-value adjusting rv + associated common variants |
| --- | --- | --- | --- | --- | --- | --- | --- | --- | --- | --- | --- | --- |
| N2-acetyllysine | Negative Control | 2 | 71934335-74868682 | Afr | 0.15 | 1.60E-16 | 0.01 | 3.30E-03 | 0.04 | 3.53E-15 | 0.01 | 3.28E-03 |
| Betaine | Negative Control | 5 | 78778375-79374979 | Afr | -0.07 | 4.19E-12 | -0.02 | 5.60E-04 | — | — | — | — |
| N-acetylglucosaminylasparagine | Negative Control | 6 | 167190769-167300620 | Amer | -0.11 | 4.61E-13 | -0.02 | 1.00E-03 | — | — | — | — |
| Cysteinylglycine | Negative Control | 16 | 87530462 - 90083514 | Amer | 0.07 | 3.91E-10 | 0.01 | 7.43E-03 | — | — | — | — |
| N-acetylarginine | Test Regions | 2 | 71934335-74810716 | Afr | 0.04 | 6.6E-22 | -0.01 | 2.02E-03 | 0.05 | 5.62E-22 | -0.01 | 2.24E-03 |
| 3-aminoisobutyrate | Test Regions | 5 | 34648620-35360348 | Amer | 0.05 | 3.04E-31 | -0.01 | 6.37E-03 | 0.05 | 1.58E-32 | -0.01 | 6.37E-03 |
| N-acetylputrescine | Test Regions | 8 | 17749979-18453195 | Amer | 0.02 | 5.99E-08 | -0.01 | 1.10E-02 | — | — | — | — |
| 5-oxoproline | Test Regions | 8 | 141782700-144007080 | Afr | -0.10 | 4.38E-25 | 0.01 | 5.90E-03 | -0.05 | 2.18E-25 | 0.01 | 3.78E-03 |
| Carnitine | Test Regions | 10 | 59301068-61869652 | Amer | 0.07 | 6.25E-17 | -0.01 | 5.49E-03 | — | — | — | — |
| Arachidonoylcholine | Test Regions | 11 | 60178829-65895768 | Amer | -0.04 | 6.2E-30 | -0.02 | 7.66E-03 | -0.04 | 2.87E-29 | -0.02 | 7.66E-03 |
| 3beta-hydroxy-5-cholestenoate | Test Regions | 11 | 107710259-111394190 | Amer | -0.04 | 5.11E-27 | -0.02 | 5.01E-05 | — | — | — | — |
| 1-methylimidazoleacetate | Test Regions | 12 | 84652-502428 | Amer | -0.04 | 1.38E-29 | -0.01 | 9.91E-03 | -0.04 | 1.74E-28 | -0.01 | 4.86E-03 |
| Ethylmalonate | Test Regions | 12 | 120613482-122018618 | Afr | -0.04 | 3.24E-15 | 0.01 | 5.04E-03 | -0.04 | 2.96E-16 | 0.01 | 7.12E-03 |
| N-acetylcarnosine | Test Regions | 13 | 95131902 - 96203528 | Afr | 0.05 | 3.26E-09 | 0.01 | 6.51E-03 | — | — | — | — |
| Propyl 4-hydroxybenzoate sulfate | Test Regions | 16 | 27060560-31617061 | Afr | 0.06 | 4.63E-34 | 0.03 | 7.75E-06 | 0.05 | 4.65E-19 | 0.03 | 7.75E-06 |
| Cys-gly, oxidized | Test Regions | 16 | 87530462 - 90083514 | Amer | 0.05 | 9.42E-34 | -0.01 | 4.27E-03 | 0.05 | 4.78E-34 | -0.01 | 4.99E-03 |

We also examined the span of associated common variants adjusted across the significant AM region. Most of the common variants are in high proximity to each other, while two variants at chromosome 2 are situated relatively far away (above 1.5 Mb) from other identified common variants but are still significantly associated with the metabolites in conditional analysis (Figure 4A).

### Admixture mapping with rare variants – test regions

For the genomic regions whose AM signals were not explained in the previous study (referred to as test regions), we first adjusted for the “associated common variants” identified through single variant analysis in combination of known genetic associations to assess how much of the admixture mapping (AM) signal could be explained and whether the results aligned with previously reported findings. In all twelve test regions, the AM signals were explained solely by adjusting for significantly associated common variants within an expanded region compared to the previous publication ^[19]^, and this pattern was consistent across both the discovery and replication batches (Table 3, and Supplementary Table S3).

To assess the impacts of rare variants in AM, we first extracted burden scores from the significant rare variant sets identified in the test regions, each associated with its respective metabolite. In regions with multiple rare variant sets, all unique gene–category pairs that were not highly correlated (r < 0.7) were jointly included as covariates in the conditional AM through their burden scores (Supplementary Figure S2). When only adjusting for the RV sets at the propyl 4-hydroxyenzoate sulfate on chromosome 16 test region, the AM signal in African ancestry became partially explained compared to the null model, but remained significant (Figure 3D, Table 3). The gap observed in (D) is attributable to missing local ancestry calls in the centromeric region, likely due to exclusion during the quality-control step. However, in the remaining regions, adjusting for rare variant sets alone resulted in little change relative to the null model (Figure 3A–C, Table 3, and Supplementary Figure S3). When adjusting for both the known common variants and the RV set burdens did not result in further change to the AM association (compared to adjusting only for common variants; Figure 3, Table 3, and Supplementary Figure S3). For the four regions with rare variant sets replicated in batch 2, the patterns were similar as those shown in batch 1 (Figure 3).

**Figure 3.**
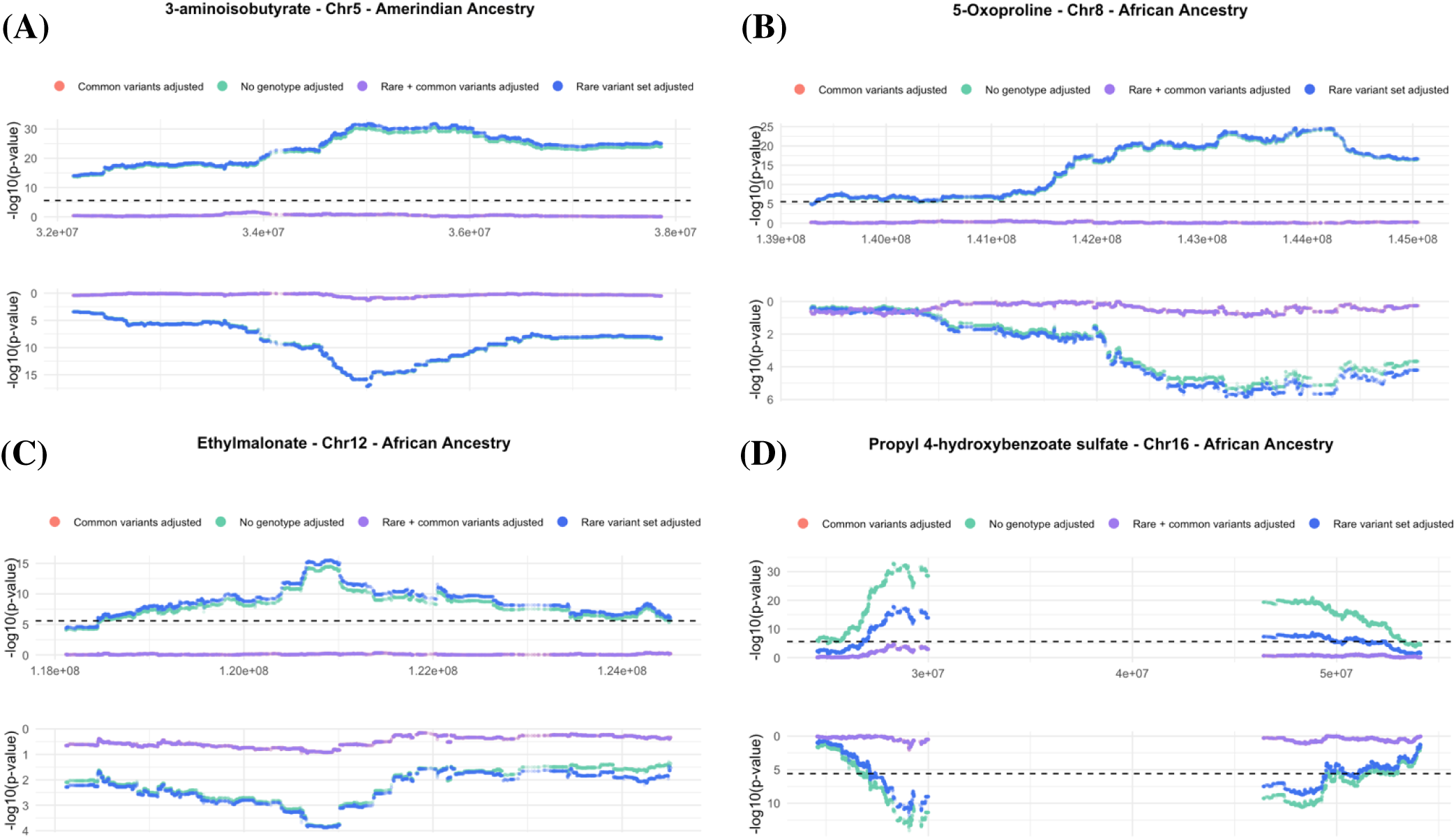
AM results in terms of -log10(P) using the (A) Amerindian local ancestry for 3-aminoisobutyrate on chromosome 5, (B) African local ancestry for 5-oxoproline on chromosome 8, (C) African local ancestry for ethylmalonate on chromosome 12, and (D) African ancestry for propyl 4-hydroxyenzoate sulfate on chromosome 16, using null model (no genotype adjusted), model adjusting common variants, model adjusting rare variant, and model adjusting both rare and common variants for discovery (upper) and replication batch (lower). The gap observed in (D) reflects the absence of local ancestry information in that region, which is near the centromere and was likely removed during quality control. The horizontal dashed black line in the upper panel denotes the AM significance threshold (P = 2.58 × 10^-6^). The rare variant sets included in the batch 2 were those that had also been replicated in batch 1.

The genomic positions of associated common variants could span a relatively wide region (1–2 Mb) on chromosomes 2 (N-acetylarginine), 8 (5-oxoproline), 10 (carnitine), and 11 (arachidonoylcholine and 3beta-hydroxy-5-cholestenoate) (Figure 4-5, Table 4). In comparison, in some regions the common variants span a narrower genomic range (Figure 4-5, Table 4).

**Figure 4.**
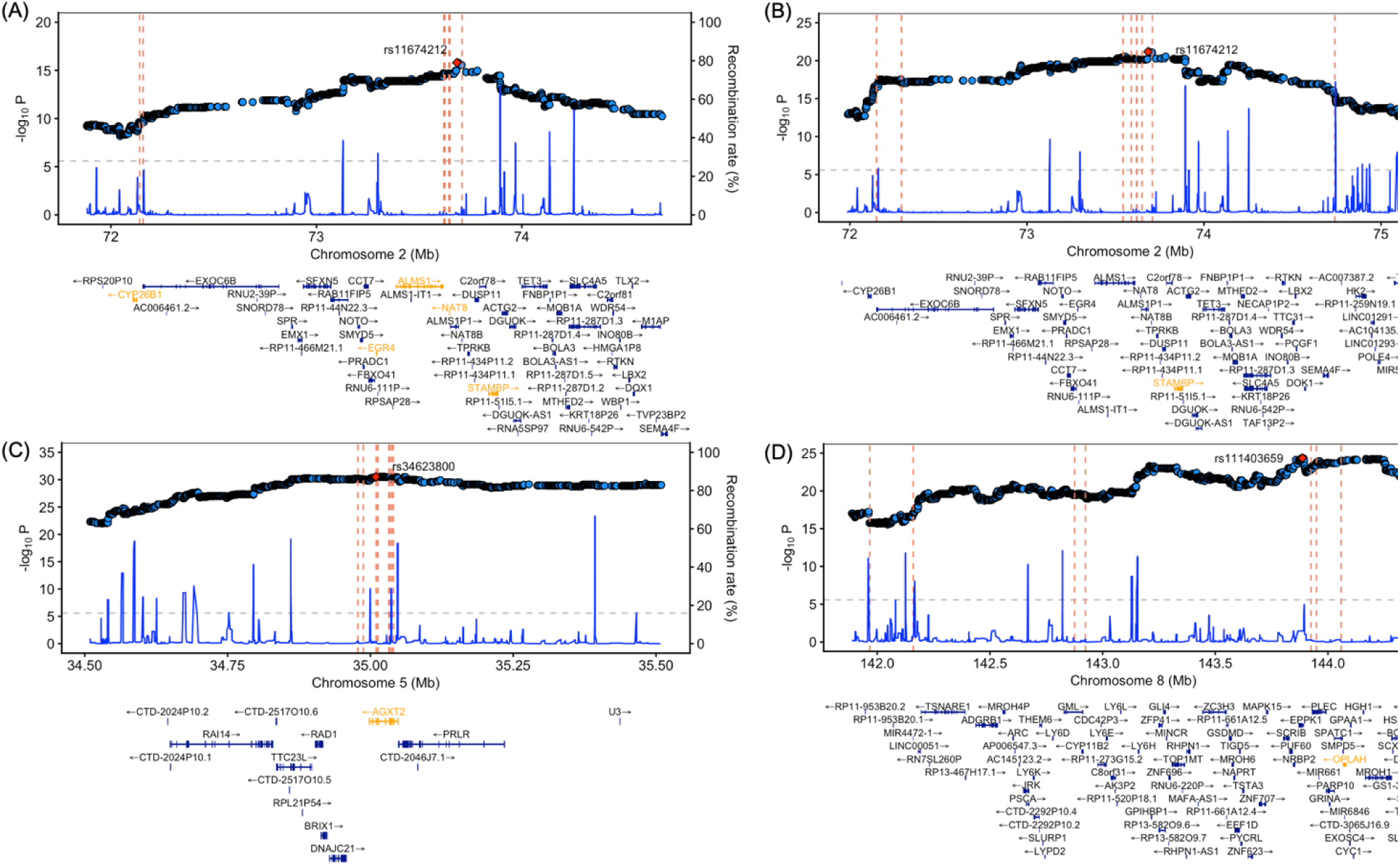
AM regional association plots of regions where significant rare variant sets were identified, showing positions of associated common variants and genes with significant rare variant sets in conditional analysis (chr 2-8). The horizontal dashed line shows AM significance threshold (P = 2.58 × 10^-6^). (A) African ancestry for N2-acetyllysine on chromosome 2, (B) African ancestry for N-acetylarginine on chromosome 2 (C) Amerindian ancestry for 3-aminoisobutyrate on chromosome 5, (D) African ancestry for 5-oxoproline on chromosome 8. Each vertical dashed line marks the position of a known common variant, while the highlighted gene denotes the genomic location and gene assignment of the rare variant sets identified in the conditional analysis.

**Figure 5.**
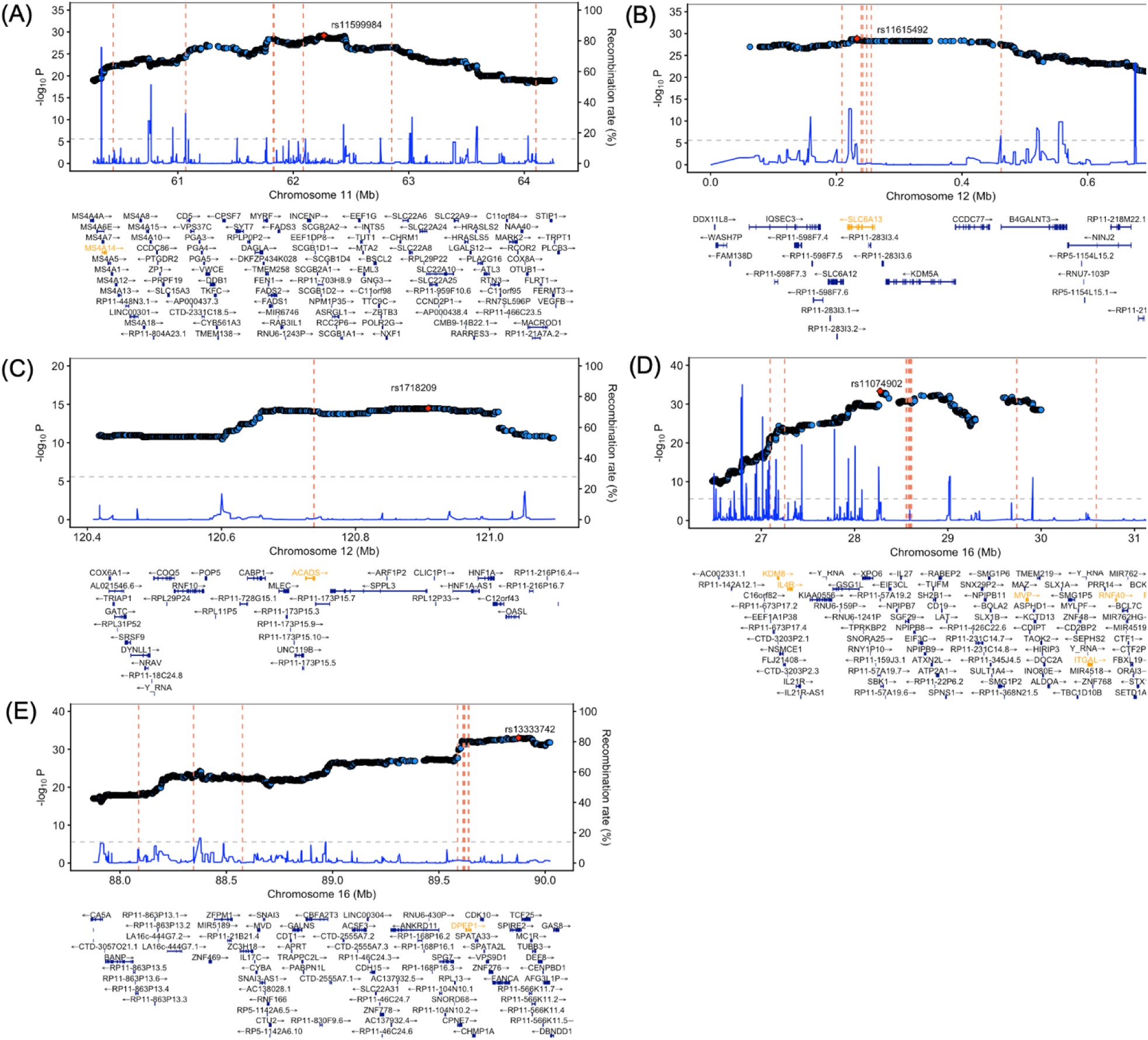
AM regional association plots showing positions of associated common variants and genes with significant rare variant sets in conditional analysis (chr 11-16). The horizontal dashed line shows AM significance threshold (P = 2.58 × 10^-6^). (A) Amerindian ancestry for arachidonoylcholine on chromosome 11, (B) Amerindian ancestry for 1-methylimidazoleacetate on chromosome 12, (C) African ancestry for ethylmalonate on chromosome 12, (D) African ancestry for propyl 4-hydroxybenzoate sulfate on chromosome 16, and (E) Amerindian ancestry for cys-gly, oxidized on chromosome 16. Each vertical dashed line marks the position of a known common variant, while the highlighted gene denotes the genomic location and gene assignment of the rare variant sets identified in the conditional analysis.

**Table 4.**
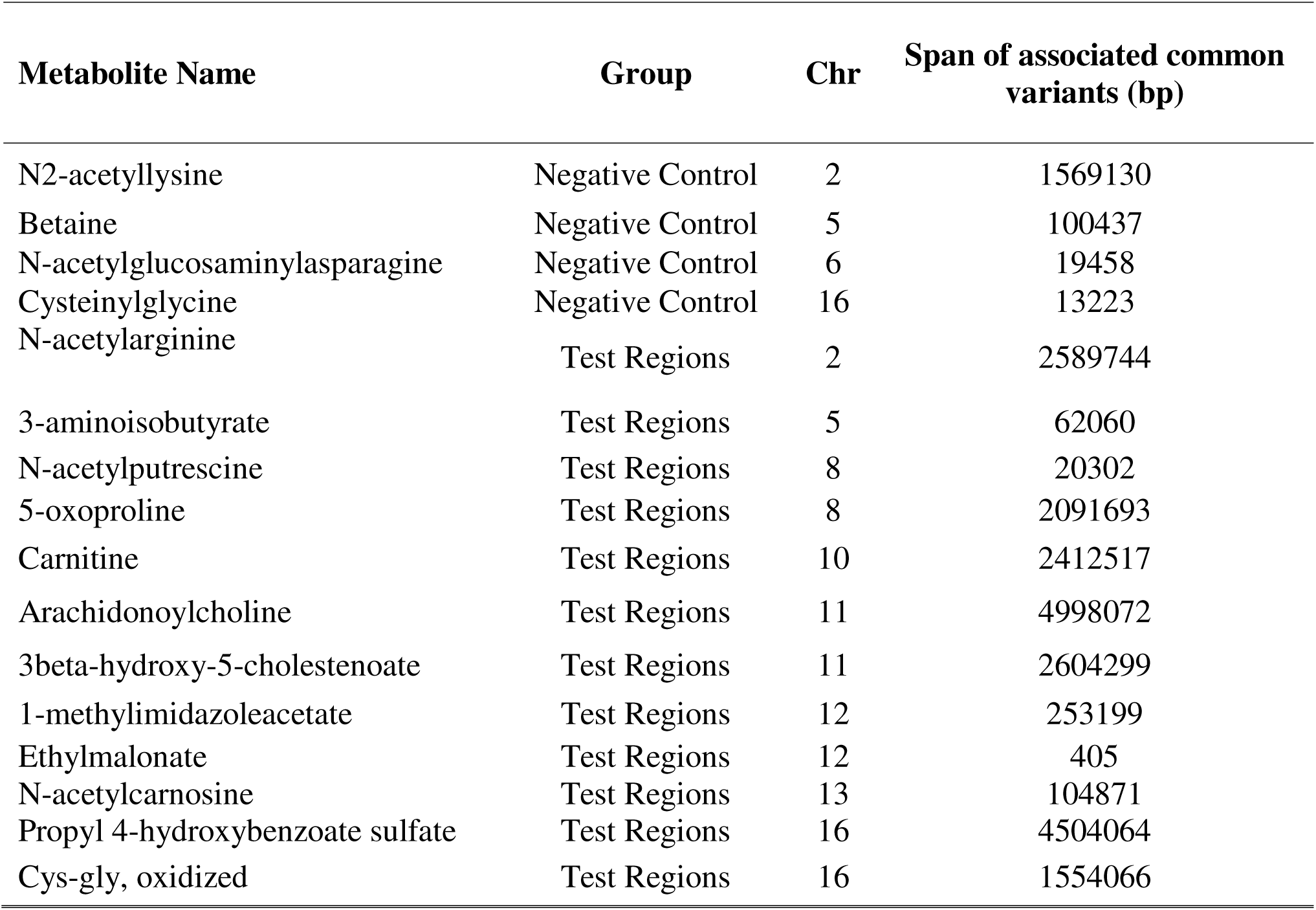
Genomic position span (in base pairs) of associated common variants across all sixteen metabolite–genomic regions.

Among all regions with rare variant sets identified in coding or non-coding conditional analysis, the mapped genes harboring these sets are very close to associated common variants (Figure 4-5).

For example, for cys-gly (oxidized) on chromosome 16, both rare and common variants are concentrated around the *DPEP1* gene, and for 3-aminoisobutyrate on chromosome 5, both rare and common variants are concentrated around the *AGXT2* ene (Figure 5E, Table 4).

## Discussion

Admixture mapping is effective in identifying ancestry-enriched regions associated with phenotypic traits by leveraging inferred local ancestries. Due to the extensive local ancestry regions in admixed populations, and unclear mechanism of local ancestry association ^[51]^, examining how AM signals can be explained is essential for discovery of specific trait-associated genetic variants. A longstanding hypothesis has been that AM associations that were not explained by common variants are driven by rare genetic variants. This work aimed to study this hypothesis.

We identified 35 metabolite-associated rare variant sets with coding or non-coding functions through gene-centric analyses for 16 selected genomic regions via the modified pipeline for rare variant associations. Further, we replicated 4 of the rare variant set associations in the replication batch. We found no significantly associated rare variant sets in some of the selected genomic regions (Table 2), and for those with rare variant sets, the genes containing those rare variants are close to the known common variants (Figure 4-5). The number of significantly associated rare variant sets dropped significantly once conditioning on the known common variants, from 104 to 35 in batch 1 and 67 to 4 in batch 2 (Table 2).

Our previous study identified genomic regions with strong AM signals, and found that these association regions could not be explained by genome-wide significant common variants suggested by COJO analysis applied over GWAS summary statistics ^[19]^. COJO first identifies the most strongly associated variant and then performs conditional association analysis adjusting for this variant to identify the next most strongly associated signal. It subsequently conditions on the newly identified SNP(s) to detect additional independent variants, repeating this iterative process until no genome-wide significant SNPs remain ^[52,53]^. The method fully explained the unexplained AM signals in the two examples provided in our previous study ^[19]^. This method is analogous to our workflow of fitting “common associated variants” in conditional AM, but differs in the methods of identifying common variants, and the results that AM signals were explained after adjustment are consistent.

Here, we defined a genomic interval within which we jointly interrogated both common and rare variants, enabling us to account for all AM signals across the four negative control regions and all twelve test regions with previously unexplained AM signals. The corresponding attenuation in AM association strength between the unconditional and conditional AM analyses was consistently replicated in batch 2 (Table 3, Supplementary Table S3).

It is expected that some of the AM signal is driven by GWAS signal. Burden scores extracted from a single rare variant set had limited ability for explaining the AM signals (Table 3, Supplementary Table S3). When there were more rare variant sets as in the test region for propyl 4-hydroxybenzoate sulfate at chromosome 16, we observed more signal reduction than those in other regions (Figure 3D). Rare variant sets identified by the modified STAAR pipeline were often located near the adjusted common variants, suggesting that the common variants and rare variant sets identified are associated with the same gene (Figure 4-5).

We note that we have a small sample size in batch 1 (n = 3,889), which reduces power to detect rare variant associations compared to biobank-scale studies. Thus, it is possible that our analysis missed rare variants sets with lower significance levels that are indeed associated with the metabolite. Admixture mapping can tag variants with ancestry-specific effects that may be missed by GWAS, making it a valuable complement for uncovering ancestry-driven regions and variants potentially under selection or associated with differential susceptibility. Therefore, examining ancestry-enriched but GWAS non-significant variants is also important – but requires replication. Note that we only identified significantly associated genetic variants, rather than studying multiple causal variants in this framework, which requires fine-mapping techniques and could be included in the future works.

To more comprehensively evaluate the role of rare variants in admixture mapping, future studies will expand the number of metabolite–genomic regions analyzed by considering variants in a wide region around AM associations and extracting burden scores. It will also be important to investigate the functional relevance of rare variant sets associated with metabolites.

## Acknowledgements

The authors thank the participants and staff of the Hispanic Community Health Study/Study of Latinos (HCHS/SOL) for their valuable contributions. A full list of investigators and study staff is available in Sorlie et al., *Annals of Epidemiology* (2010 Aug; 20:642–649) and on the study website (http://www.cscc.unc.edu/hchs/). HCHS/SOL is a collaborative study supported by contracts from the National Heart, Lung, and Blood Institute (NHLBI) awarded to the University of North Carolina, University of Miami, Albert Einstein College of Medicine, University of Illinois at Chicago, and San Diego State University. Additional support was provided through interagency transfers to NHLBI from the National Institute on Minority Health and Health Disparities; the National Institute on Deafness and Other Communication Disorders; the National Institute of Dental and Craniofacial Research; the National Institute of Diabetes and Digestive and Kidney Diseases; the National Institute of Neurological Disorders and Stroke; and the NIH Office of Dietary Supplements. Support for metabolomics data was graciously provided by the JLH Foundation (Houston, Texas) and by NHLBI grant R01HL141824. This study was supported by National Human Genome Research Institute grants R56HG013163 and R01HG013163.

## Ethics statement

The HCHS/SOL study received ethical approval from institutional review boards (IRBs) at each study site, and all participants provided written informed consent. Oversight for the Data Coordinating Center was provided by the Non-Biomedical IRB at the University of North Carolina at Chapel Hill. The IRBs that reviewed and approved the study include: UNC-Chapel Hill’s Non-Biomedical IRB (Chapel Hill, NC), the Einstein IRB at Albert Einstein College of Medicine (Bronx, NY), the IRB at the University of Illinois at Chicago (Chicago, IL), the Human Subject Research Office at the University of Miami (Miami, FL), and the IRB at San Diego State University (San Diego, CA). All research activities involving HCHS/SOL data and biospecimens were conducted in compliance with applicable ethical regulations for human subjects research. This work was additionally reviewed and approved by the Committee on Clinical Investigations at Beth Israel Deaconess Medical Center.

## Code availability statement

The code for adapted STAAR pipeline, conditional AM, and figures and tables are available at the github repository: https://github.com/xc448/HCHS_SOL_Rare_Variants

## Data availability statement

The HCHS/SOL fully supports data sharing for HCHS/SOL–approved manuscript proposals with outside investigators. All data sharing is conducted in accordance with HCHS/SOL study and NIH policies and governed by a Data and Materials Distribution Agreement (DMDA) between UNC and the external institution, ensuring the confidentiality and privacy of HCHS/SOL participants and their families. Alternatively, de-identified HCHS/SOL data are publicly available at BioLINCC and dbGaP for the subset of the study cohort who authorized general use of their data at the time of informed consent. HCHS/SOL WGS data are available by application to dbGaP according to the study-specific accession “phs001395”. Phenotype data are also available from dbGaP according to the study-specific accession “phs000810”.

## Author contribution statement

TS planned and supervised the study. XC prepared the modified STAAR pipeline, and performed all, computational and statistical analyses, figure creations, and manuscript writing. All authors contributed to manuscript editing.

