## Supplementary Materials for "Explaining the unexplained admixture mapping signals via rare variants: the HCHS/SOL"

### Supplementary Figures

#### Supplementary Figure S1: Genetic PCs computed from two metabolomic batched combined, grouped by self-identified backgrounds

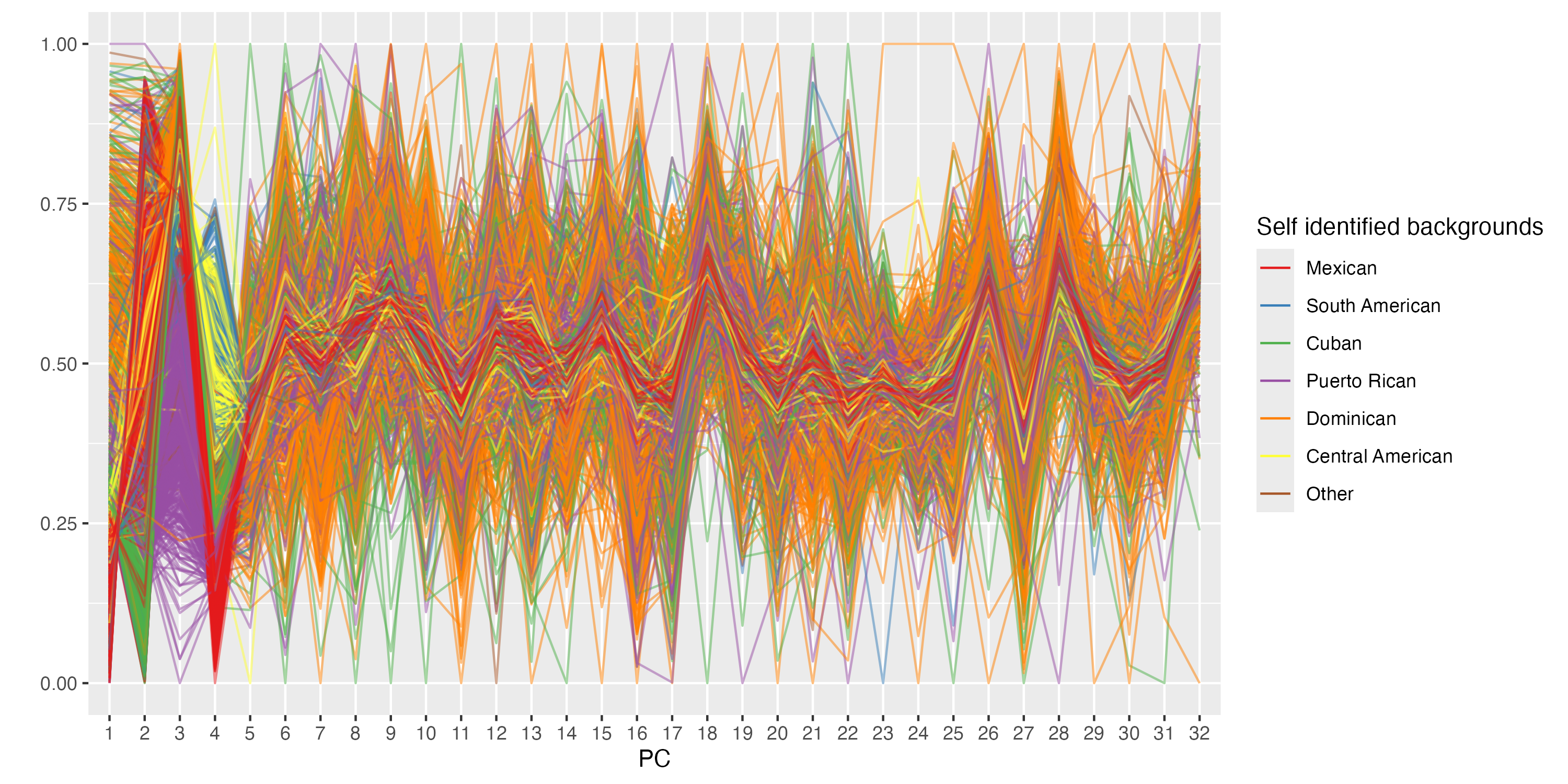

**(A)**

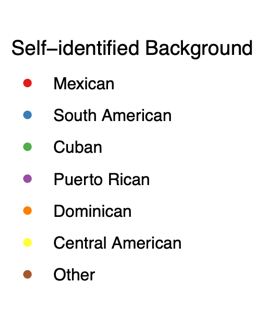

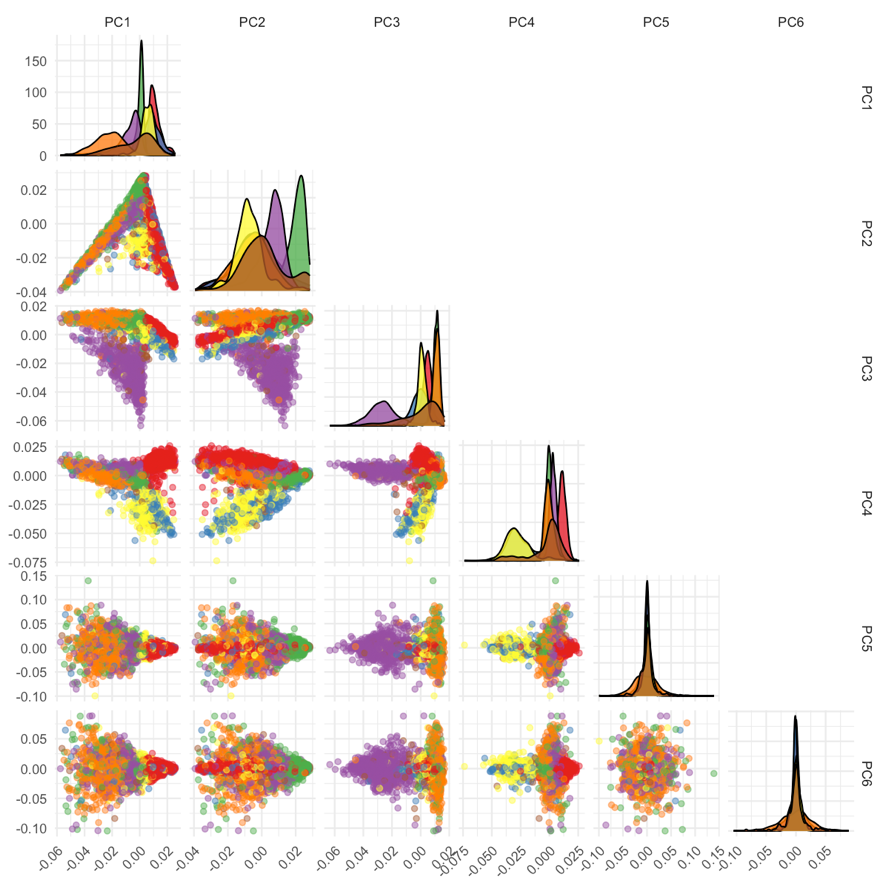

**(B)**

Principal components (PCs) of all individuals from the metabolomic batches excluding outliers with elevated East Asian ancestry, color-coded by self-identified backgrounds. The “Other” category includes participants who reported multiple or unspecified / missing background information. (A) Plot of parallel coordinates for the first 32 PCs. The 32 parallel vertical lines of equal length correspond to the first 32 PCs. Each individual is represented by a set of line segments connecting his or her PC values. (B) Pair-wise PC values from PC1 to PC6, each dot representing an individual, colored by self-identified background.

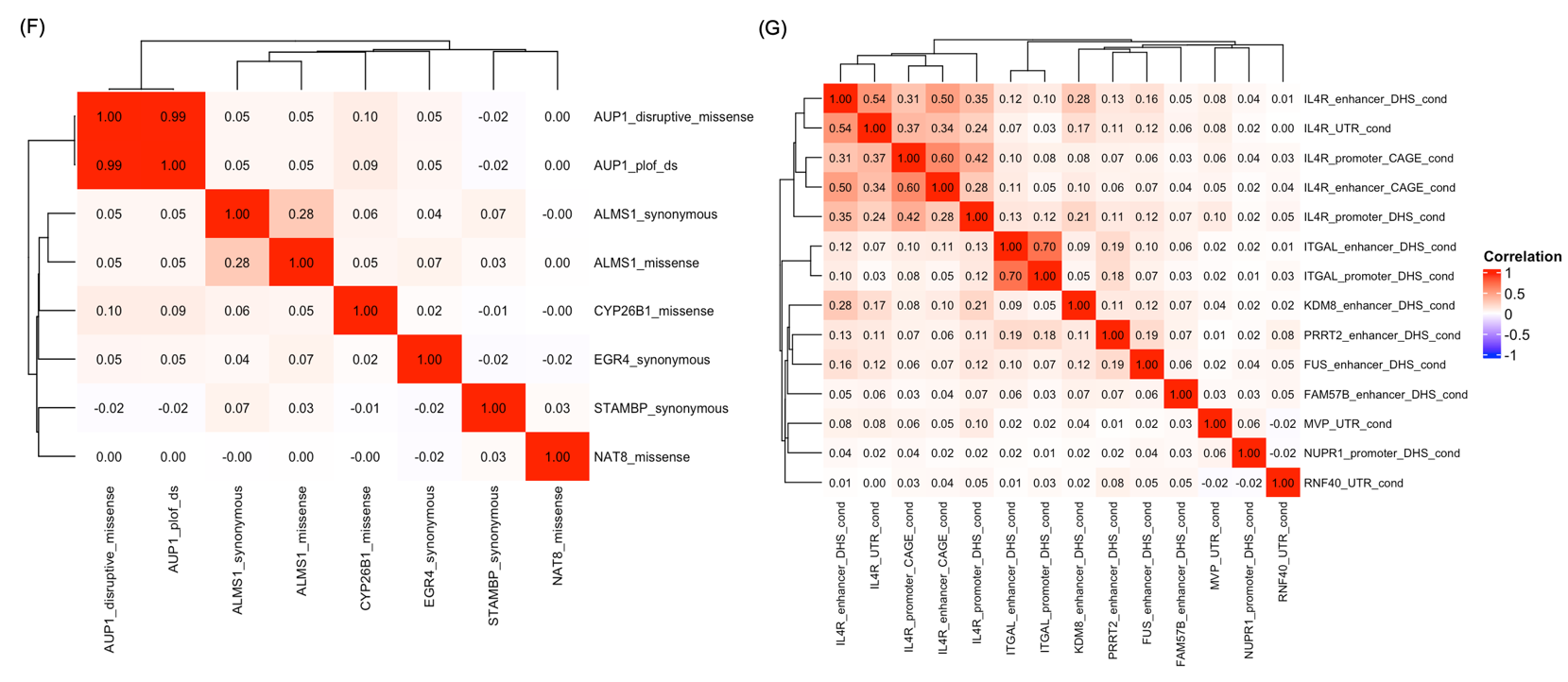

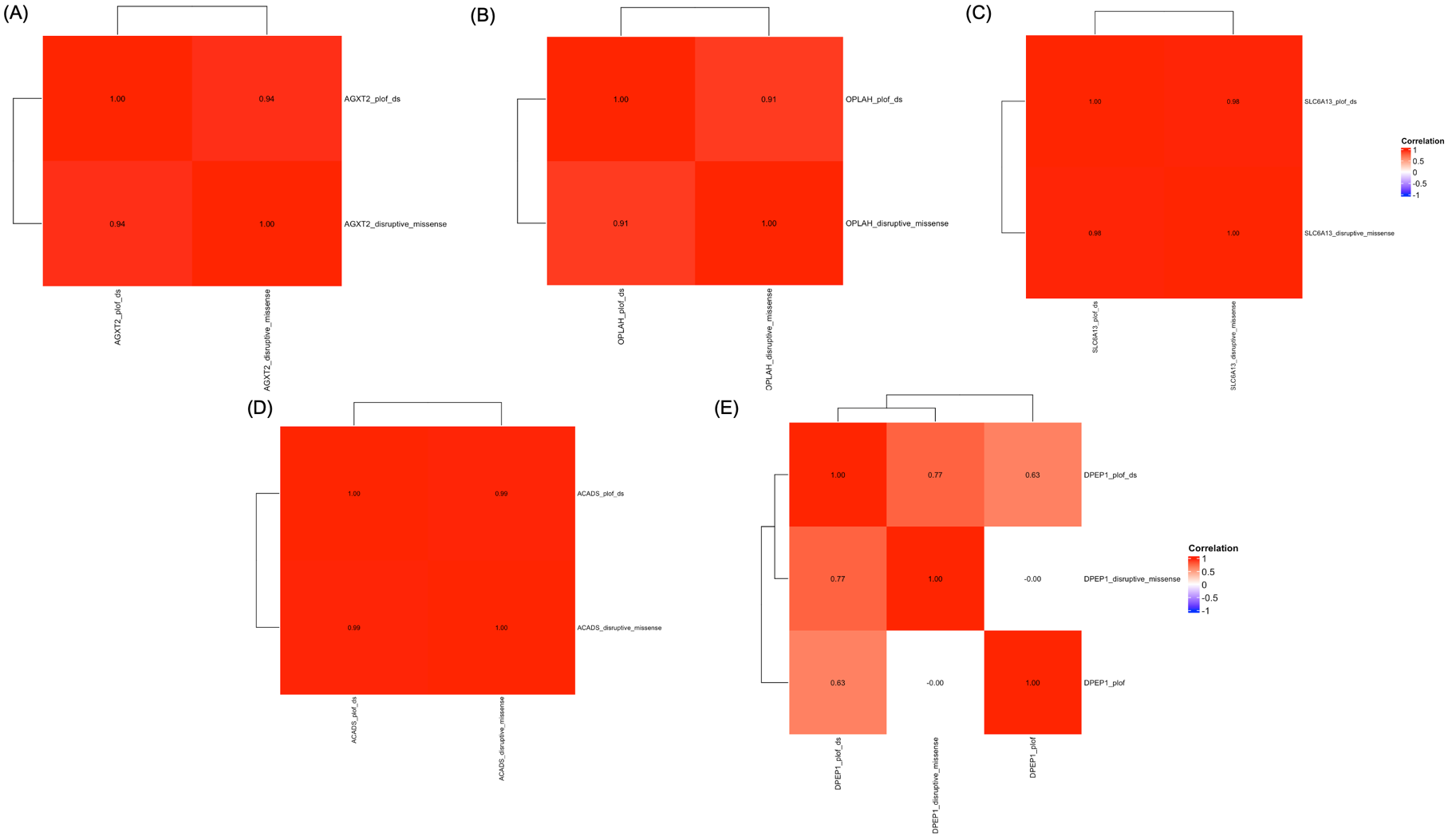

#### Supplementary Figure S2: Correlation heatmaps of rare variant sets identified in the adapted STAAR pipeline across genomic regions

Heatmaps depicting Pearson correlations among significant rare variant (RV) sets identified in regions of interest in the discovery batch: (A) RV sets at chr5 (*AGXT2*) associated with 3-aminoisobutyrate; (B) RV sets at chr8 (*OPLAH*) associated with 5-oxoproline; (C) RV sets at chr12 (*SLC6A13*) associated with 1-methylimidazoleacetate; (D) RV sets at chr12 (*ACADS*) associated with ethylmalonate; (E) RV sets at chr16 (*DPEP1*) associated with cysteinylglycine (oxidized); (F) RV sets at chr2 mapped to multiple genes associated with N2-acetyllysine; and (G) RV sets at chr16 mapped to multiple genes associated with propyl 4-hydroxybenzoate sulfate.

#### Supplementary Figure S3: AM results showing batch 1 and batch 2 results with rare variant sets only identified in batch 1

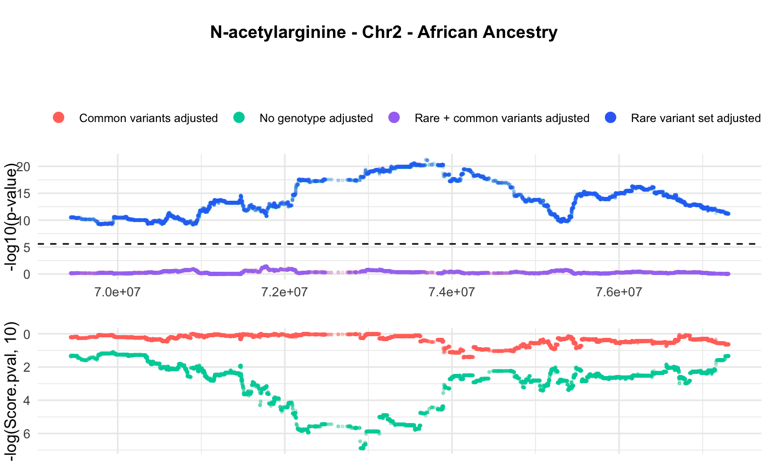

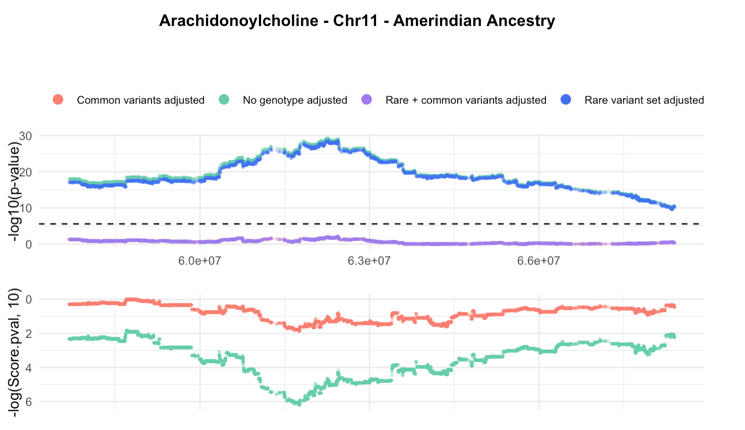

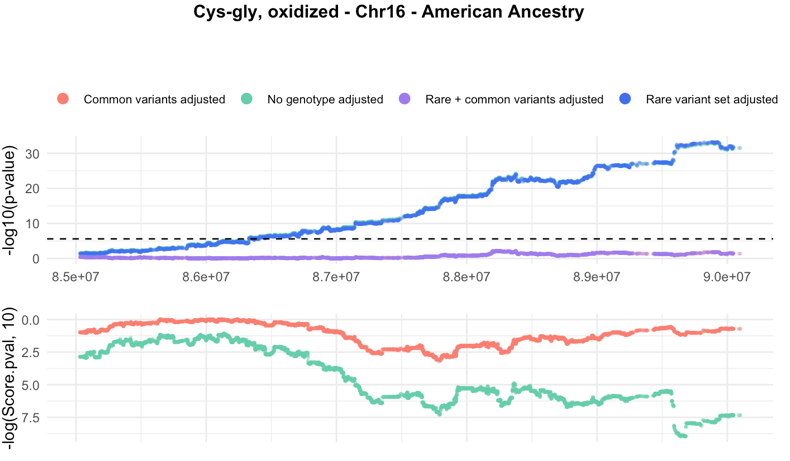

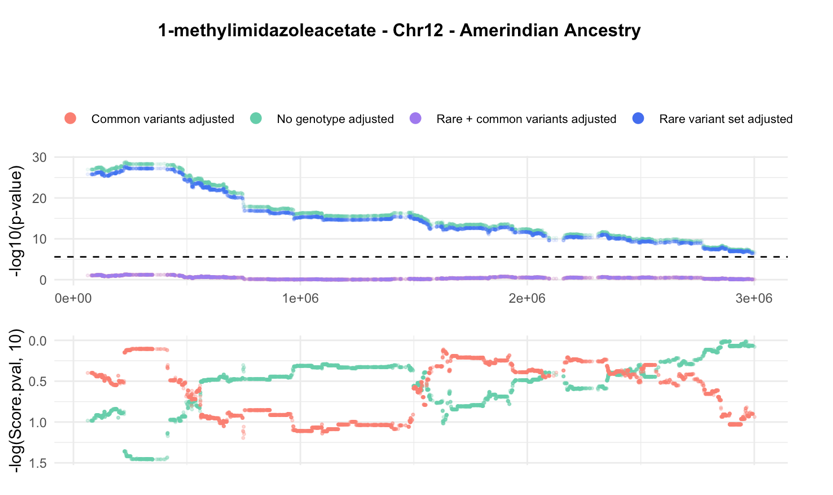

**(C)**

**(D)**

**(A)**

**(B)**

AM results in terms of -log10(P) using the driving local ancestries for four test region traits: (A) N-acetylarginine on chromosome 2 driven by African ancestry, (B) arachidonoylcholine on chromosome 11 driven by Amerindian ancestry, (C) 1-methylimidazoleacetate on chromosome 12 driven by Amerindian ancestry and (D) cys-gly, oxidized on chromosome 16 driven by Amerindian ancestry, using null model (no genotype adjusted), model adjusting common variants, model adjusting rare variant, and model adjusting both rare and common variants for discovery (upper) and replication batch (lower). The horizontal dashed black line in the upper panel denotes the AM significance threshold (P = 2.58 $\times$10^-6^). The rare variant sets included in the batch 1not replicated in batch 2 therefore not included in the batch 2 results.

### Supplementary Tables

#### Supplementary Table S1: Characteristics of participants included in admixture mapping

|  | **Discovery Batch (N=3842)** | **Replication Batch (N=1465)** | **Overall (N=5307)** |
| --- | --- | --- | --- |
| **Age (years)** |  |  |  |
| Mean (SD) | 45.9 (13.8) | 52.5 (10.9) | 47.7 (13.4) |
| **BMI (kg/m^2^)** |  |  |  |
| Mean (SD) | 29.8 (6.05) | 30.2 (6.10) | 29.9 (6.06) |
| **eGFR (mL/min/1.73m^2^)** |  |  |  |
| Mean (SD) | 109 (26.5) | 105 (25.8) | 108 (26.3) |
| **Gender** |  |  |  |
| Female | 2194 (57.1%) | 928 (63.3%) | 3122 (58.8%) |
| Male | 1648 (42.9%) | 537 (36.7%) | 2185 (41.2%) |
| **Recruitment Center** |  |  |  |
| Bronx | 1016 (26.4%) | 413 (28.2%) | 1429 (26.9%) |
| Chicago | 874 (22.7%) | 312 (21.3%) | 1186 (22.3%) |
| Miami | 1086 (28.3%) | 463 (31.6%) | 1549 (29.2%) |
| San Diego | 866 (22.5%) | 277 (18.9%) | 1143 (21.5%) |
| **Self-reported Background** |  |  |  |
| Mexican | 1390 (36.2%) | 411 (28.1%) | 1801 (33.9%) |
| South American | 225 (5.9%) | 96 (6.6%) | 321 (6.0%) |
| Cuban | 652 (17.0%) | 330 (22.5%) | 982 (18.5%) |
| Puerto Rican | 685 (17.8%) | 272 (18.6%) | 957 (18.0%) |
| Dominican | 378 (9.8%) | 205 (14.0%) | 583 (11.0%) |
| Central American | 386 (10.0%) | 133 (9.1%) | 519 (9.8%) |
| Other | 126 (3.3%) | 18 (1.2%) | 144 (2.7%) |
| **Hypertension** | 1103 (28.7%) | 593 (40.5%) | 1696 (32.0%) |
| **Diabetes** | 718 (18.7%) | 418 (28.5%) | 1136 (21.4%) |

Participant characteristics from either the discovery batch 1 or the replication batch 2 included in admixture mapping. Categorical variables are reported with proportion per level and continuous variables are reported with mean and standard deviation (SD), eGFR: estimated glomerular filtration rate.

| Metabolite Name  Supplementary Table S2: Summary of significant rare variant sets from all regions, both coding and non-coding gene-centric analysis in replication batch 2 | | Type | | Chr | | Region  (bp) | | Driving Ancestry | | # previously reported independent variants | | # significant individual variants, conditional P < 0.001, pruned | | # significant RV sets, coding (unconditional,  P < 2.71E-05) | | # unique significant RV sets, coding (conditional, P < 0.001) | # significant RV sets, non-coding (unconditional, P < 1.47E-05) | | # significant RV sets, non-coding (conditional, P < 0.001) |
| --- | --- | --- | --- | --- | --- | --- | --- | --- | --- | --- | --- | --- | --- | --- | --- | --- | --- | --- | --- |
| N2-acetyllysine | | Negative Control | | 2 | | 71934335-74868682 | | Afr | | 1 | | 5 | | 1 | | 0 | 2 | | 0 |
| Betaine | | Negative Control | | 5 | | 78778375-79374979 | | Afr | | 2 | | 0 | | 0 | | -- | 0 | | -- |
| N-acetylglucosaminylasparagine | | Negative Control | | 6 | | 167190769-167300620 | | Amer | | 2 | | 0 | | 0 | | -- | 0 | | -- |
| Cysteinylglycine | | Negative Control | | 16 | | 87530462 -90083514 | | Amer | | 1 | | 2 | | 2 | | 3 (0) | 9 | | 10 (0) |
| N-acetylarginine | | Test Region | | 2 | | 71934335-74810716 | | Afr | | 3 | | 7 | | 1 | | 0 | 0 | | -- |
| 3-aminoisobutyrate | | Test Region | | 5 | | 34648620- 35360348 | | Amer | | 5 | | 5 | | 5 | | 2* (1) | 0 | | -- |
| N-acetylputrescine | | Test Region | | 8 | | 17749979- 18453195 | | Amer | | 3 | | 3 | | 1 | | 0 | 1 | | 0 |
| 5-oxoproline | | Test Region | | 8 | | 141782700-144007080 | | Afr | | 2 | | 8 | | 0 | | 1 (1) | 0 | | -- |
| Carnitine | | Test Region | | 10 | | 59301068-61869652 | | Amer | | 3 | | 3 | | 0 | | 0 | 0 | | 0 |
| Arachidonoylcholine | | Test Region | | 11 | | 60178829- 65895768 | | Amer | | 1 | | 8 | | 2 | | 0 | 0 | | -- |
| 3beta-hydroxy-5-cholestenoate | | Test Region | | 11 | | 120613482- 122018618 | | Amer | | 2 | | 3 | | 0 | | -- | 0 | | -- |
| 1-methylimidazoleacetate | | Test Region | | 12 | | 84652-502428 | | Amer | | 2 | | 4 | | 0 | | -- | 0 | | -- |
| Ethylmalonate | | Test Region | | 12 | | 120613482- 122018618 | | Afr | | 2 | | 0 | | 4 | | 2*(1) | 1 | | -- |
| N-acetylcarnosine | | Test Region | | 13 | | 95131902 -96203528 | | Afr | | 3 | | 1 | | 1 | | -- | 0 | | -- |
| Propyl 4-hydroxybenzoate sulfate | | Test Region | | 16 | | 27060560- 31617061 | | Afr | | 3 | | 15 | | 10 | | 0 | 26 | | 3 (3) |
| Cys-gly, oxidized | | Test Region | | 16 | | 87530462 -90083514 | | Amer | | 5 | | 3 | | 1 | | 0 | 1 | | 0 |
| Number of significant single common variants and rare variant sets identified using the modified STAAR pipeline in batch 2, both conditional and unconditional on the “associated common variants” for each trait and genomic region, which were generated by combining the variants from the 6^th^ and 7^th^ column in the table together. Afr: African ancestry, Amer: Amerindian ancestry, RV: rare variants. Entries marked with “*” indicate that some identified rare variant sets shared identical variant composition and burden scores; the counts shown therefore represent the deduplicated number. Entries marked with “()” indicate the total number of rare variant sets that are replicated in batch 1 as shown in Table S4. |  | |  | |  | |  | |  | |  | |  | |  | |  |  | |

#### Supplementary Table S3: Replicated batch 2 rare variant set test results

| **Gene Name** | **Chr** | **Metabolite** | **Group** | **Category** | **#SNV** | **cMAC** | **STAAR-S**  **(1,25)** | **STAAR-S**  **(1,1)** | **STAAR-B**  **(1,25)** | **STAAR-B**  **(1,1)** | **STAAR-A**  **(1,25)** | **STAAR-A**  **(1,1)** | **ACAT-O** | **STAAR-O** |
| --- | --- | --- | --- | --- | --- | --- | --- | --- | --- | --- | --- | --- | --- | --- |
| AGXT2 | 5 | 3-aminoisobutyrate | Test Regions | plof_ds, disruptive missense | 6 | 12 | 2.99E-06 | 3.36E-06 | 1.80E-10 | 1.98E-10 | 1.80E-10 | 1.98E-10 | 3.68E-10 | 2.83E-10 |
| OPLAH | 8 | 5-oxoproline | Test Regions | plof_ds | 10 | 15 | 2.37E-03 | 2.44E-03 | 1.77E-04 | 1.73E-04 | 1.77E-04 | 1.73E-04 | 2.54E-04 | 2.54E-04 |
| ACADS | 12 | Ethylmalonate | Test Regions | plof_ds, disruptive missense | 20 | 71 | 1.12E-04 | 1.28E-04 | 1.07E-05 | 8.90E-06 | 7.01E-04 | 6.77E-04 | 2.59E-05 | 2.66E-05 |
| FAM57B | 16 | Propyl 4-hydroxybenzoate sulfate | Test Regions | enhancer_DHS | 45 | 142 | 7.09E-02 | 9.64E-02 | 7.18E-04 | 8.88E-04 | 1.11E-02 | 1.48E-02 | 1.70E-03 | 2.22E-03 |
| PRRT2 | 16 | Propyl 4-hydroxybenzoate sulfate | Test Regions | enhancer_DHS | 180 | 526.17 | 1.72E-04 | 5.29E-04 | 4.91E-01 | 8.64E-01 | 2.34E-02 | 2.34E-02 | 3.76E-04 | 7.72E-04 |
| RNF40 | 16 | Propyl 4-hydroxybenzoate sulfate | Test Regions | UTR | 17 | 57 | 2.89E-04 | 2.59E-04 | 4.80E-02 | 3.42E-02 | 2.04E-04 | 2.00E-04 | 3.42E-04 | 3.47E-04 |

Significant rare variant sets in batch 2 that replicated those identified in batch 1 based on functional category, along with their associated coding or non-coding functional annotations as determined by the modified STAAR pipeline for each metabolite and genomic region, with analyses conditioned on the corresponding associated common variants. The reported p-values are omnibus p-values derived from multiple rare variant tests (SKAT, Burden, and ACAT), with the final integrated p-value summarized as STAAR-O. STAAR-S: the omnibus p-value for all SKAT tests under the specified Beta distribution. STAAR-B: the omnibus p-value for all Burden tests under the specified Beta distribution. STAAR-A: the omnibus p-value for all ACAT-V tests under the specified Beta distribution. ACAT-O: the omnibus p-value for all aggregated tests including the SKAT, Burden, and ACAT-V tests across two Beta distributions using Cauchy method. STAAR-O: the omnibus p-value for all aggregated tests including the SKAT, Burden, and ACAT-V tests weighted by annotation using Cauchy method.

#### Supplementary Table S4: Admixture mapping results for replication batch 2

| Metabolite Name | Type | Chr | Region (bp) | Driving Ancestry | AM lead variant effect size estimates null model | AM lead variant p-value null model | AM lead variant effect size estimates cond. associated common variants | AM lead variant p-value cond. associated common variants | AM lead variant Est. adjusting rv set only | AM lead variant p-value adjusting rv set only | AM lead variant Est. adjusting rare + associated common variants | AM lead variant p-value adjusting rv + associated common variants |
| --- | --- | --- | --- | --- | --- | --- | --- | --- | --- | --- | --- | --- |
| N2-acetyllysine | Negative Control | 2 | 71934335-74868682 | Afr | 0.16 | 4.30E-06 | 0.09 | 0.01 | – | – | – | – |
| Betaine | Negative Control | 5 | 78778375-79374979 | Afr | -0.04 | 1.23E-03 | 0.03 | 6.95E-03 | – | – | – | – |
| N-acetylglucosaminylasparagine | Negative Control | 6 | 167190769-167300620 | Amer | -0.08 | 1.00E-06 | -0.05 | 6.67E-03 | – | – | – | – |
| Cysteinylglycine | Negative Control | 16 | 87530462 -90083514 | Amer | 0.08 | 4.36E-05 | -0.05 | 0.02 | – | – | – | – |
| N-acetylarginine | Test Regions | 2 | 71934335-74810716 | Afr | 0.15 | 1.30E-07 | 0.11 | 1.95E-05 | – | – | – | – |
| 3-aminoisobutyrate | Test Regions | 5 | 34648620-35360348 | Afr | 0.26 | 7.65E-18 | -0.08 | 1.90E-03 | 0.25 | 7.51E-18 | -0.08 | 2.18E-03 |
| N-acetylputrescine | Test Regions | 8 | 17749979-18453195 | Amer | -0.03 | 0.05 | -0.04 | 0.02 | – | – | – | – |
| 5-oxoproline | Test Regions | 8 | 141782700-144007080 | Afr | -0.06 | 3.39E-06 | 0.04 | 4.61E-04 | -0.06 | 1.45E-06 | 0.04 | 6.45E-04 |
| Carnitine | Test Regions | 10 | 59301068-61869652 | Amer | 0.03 | 5.68E-05 | 0.02 | 3.01E-03 | – | – | – | – |
| Arachidonoylcholine | Test Regions | 11 | 60178829-65895768 | Amer | -0.16 | 6.26E-07 | 0.10 | 3.45E-03 | – | – | – | – |
| 3beta-hydroxy-5-cholestenoate | Test Regions | 11 | 107710259-111394190 | Amer | -0.14 | 6.45E-09 | -0.07 | 2.15E-03 | – | – | – | – |
| 1-methylimidazoleacetate | Test Regions | 12 | 84652-502428 | Amer | -0.05 | 0.01 | -0.05 | 0.03 | – | – | – | – |
| Ethylmalonate | Test Regions | 12 | 120613482-122018618 | Afr | -0.28 | 1.40E-04 | 0.21 | 4.61E-03 | -0.27 | 1.31E-04 | 0.22 | 2.68E-03 |
| N-acetylcarnosine | Test Regions | 13 | 95131902 -96203528 | Afr | 0.09 | 7.93E-04 | 0.07 | 9.03E-03 | – | – | – | – |
| Propyl 4-hydroxybenzoate sulfate | Test Regions | 16 | 27060560-31617061 | Afr | 0.29 | 6.40E-15 | -0.12 | 2.69E-04 | 0.25 | 7.35E-11 | -0.12 | 1.99E-04 |
| Cys-gly, oxidized | Test Regions | 16 | 87530462 -90083514 | Amer | 0.12 | 1.12E-09 | 0.07 | 6.02E-04 | – | – | – | – |

AM results for all 16 metabolite–genomic region pairs are shown using the replication batch (batch 2) data, including effect size estimates and p-values of the lead AM variant under four models: the null model (no genotype adjustment), adjustment for known common variants only, adjustment for rare variant sets using burden scores, and adjustment for both common and rare variants (if a rare variant set was identified and previously replicated in batch 1).
